# From hours to days: prolonging the functional viability of acute brain slices to >36 h

**DOI:** 10.64898/2026.10.23.760385

**Authors:** Elizaveta Vylekzhanina, Luca Habelt, Ashlyn Creamer, Christian Cameron de Abos y Padilla, Ilka Diester, Michael Nießing, Peter Hochstrate, Philippe Coulon

**Affiliations:** Faculty of Biology, Institute of Biology III, Workgroup Cellular Neurophysiology, University of Freiburg, Germany; Faculty of Biology, Institute of Biology III, Chair of Optophysiology, University of Freiburg, Germany; Intelligent Machine-Brain Interfacing Technology//BrainLinks-BrainTools Center of Excellence, University of Freiburg, Germany; Department of Neuronal Cell Biology, Medical University of Vienna, Vienna, Austria; Institute of Neurobiology, Heinrich-Heine-University Düsseldorf, Germany

**Keywords:** mouse, 3R, acute brain slice, patch-clamp, electrophysiology, medial prefrontal cortex, motor cortex, globus pallidus, thalamic reticular nucleus, medicinal leech, artificial cerebrospinal fluid, pyruvate, malate, extended viability brain slice

## Abstract

We have developed an experimental protocol for acute brain slices of juvenile and adult mice which, in our hands, enables whole-cell patch-clamp experiments on healthy neurons with intact electrophysiological characteristics for more than 36 hours after preparation. This drastically expands the conventional viability window of brain slices of 8-12 hours, using only standard laboratory equipment and procedures. We successfully applied this protocol to evaluate neurons of the medial prefrontal cortex (mPFC), motor cortex (M1/M2), globus pallidus (GP), and thalamic reticular nucleus (TRN) of mice aged P11 to ∼2 years postnatal, demonstrating the robustness of our method. By prolonging the usability of the preparation, this procedure a) reduces the number of animals used per study, b) yields healthy brain slices also from adult animals, c) allows prolonged experimental procedures, and d) reduces material cost and preparation effort for the experimenter.

## 1. Introduction

There is consensus among scientists that the welfare of experimental animals, must be a central aim, as reflected by the "3Rs principle": 1) the necessity to *refine* experiments to minimize animal suffering and decrease the severity of procedures, 2) *reduce* the number of animals used per study and in each experiment to the minimum required to obtain meaningful results, and 3) to *replace* experimentation on higher animals by alternative methods [1,2]. However, neuroscientists seem to struggle to reduce animal use, often simply due to lack of validated alternatives. Although model organisms, such as the medicinal leech or other invertebrates, were established over 150 years ago, they could not replace research on mammalian brains, be it for *in vivo* assessment of neuronal activity during behaviour, or acute brain slice preparations, organotypical slice cultures, or primary neuronal cell cultures.

Experimental neuroscience benefits from both a tight control of physiological experimental parameters and intact system interactions. A typical model organism in neurobiological research is the medicinal leech, in which identifiable neurons and glial cells within distinct compartments (segmental ganglia) can be investigated on a cellular and sub-cellular level, while retaining the anatomy and structural integrity of the neuronal network [3–6] (see Fig. 1). An enormous wealth of literature of the study of the leech nervous system has evolved over the last century. Classic examples were the study of neuronal responses to stimulation of the leech skin in semi-intact preparations [7,8], involving both entire segments of the skin as well as intact primary sensory receptors, and complex processes such as the neuronal circuitry and activity responsible for swimming behaviour [9].

**Fig. 1:**
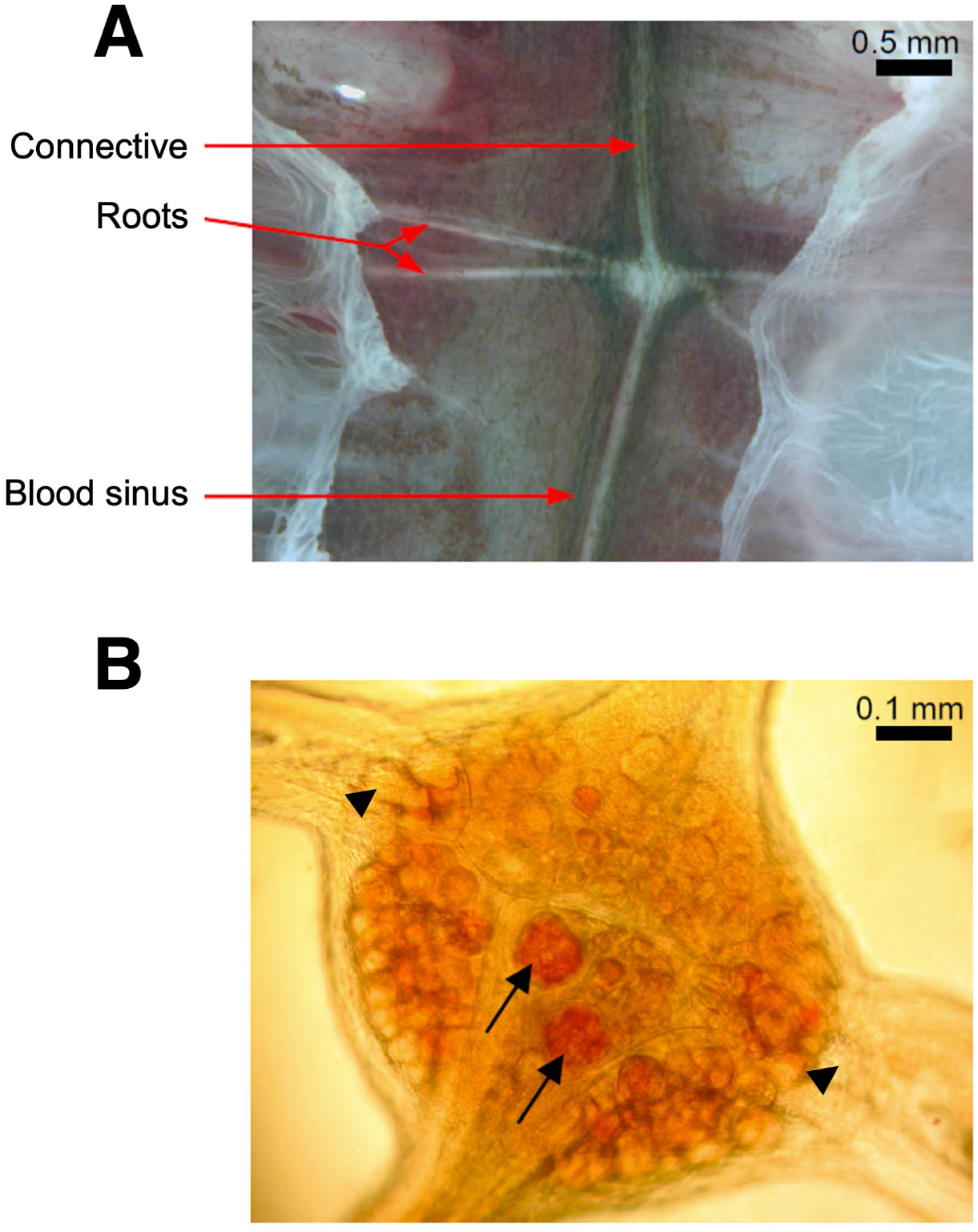
Leech segmental ganglia. **A)** A single leech segmental ganglion *in situ*. The ganglion is connected to the anterior and posterior ganglion by bundles of nerve fibres (connectives). Nerve roots connect to the innervated body segment. The ganglion chain is contained within a large blood vessel (ventral blood sinus). **B)** Neuron cell bodies in the ventral part of a segmental ganglion. Arrows mark the two serotonergic Retzius neurons which were stained with neutral red. Arrowheads mark presumed Leydig neurons.

The mammalian nervous system became more accessible with the advent of acute brain slice preparations, first developed by Li and McIlwain in 1957 [10], which opened up another compromise between intact system interactions and tight control of experimental parameters: Acute brain slices leave a part of the neuronal network intact, while allowing to control experimental parameters such as temperature, ion composition, and pH, as well as the defined application of drugs, eliminating perturbations from anaesthetics or muscle relaxants, and allowing neurons to be located, identified, accessed, and manipulated [11,12]. In some aspects, slice preparations combine the benefits and the experimental characteristics of both whole brain *in situ* and cell cultures *in vitro* [13]. The brain slice preparation is therefore considered one of the landmarks in the discipline of neuroscience [14] and today, brain slices are frequently used for electrophysiological experiments and will surely continue to be a viable tool for the foreseeable future [11].

The composition of the artificial cerebrospinal fluid used during the slicing procedure [15], for holding or incubation of acute brain slices, and during recording [16,17] has always been of particular interest, as has the slicing method, tissue oxygenation, and the design of laminar flow systems to perfuse the experimental chambers [13,17,18]. Criteria used to assess the viability or healthiness of brain slices have been refined with experimental progress [17,19]. Previous approaches were often based on the idea that an optimal environment for brain slices could be ensured by mimicking the *in vivo* situation as closely as possible [13]. However, during the slicing procedure, excitotoxicity is an undesired, albeit physiological side-effect. This may be suppressed by replacing Na^+^ with N-methyl-D-glucamine (NMDG) in the artificial cerebrospinal fluid (recovery-ACSF, [16]), which we use immediately after decapitation until the slicing procedure is complete. A further challenge is posed by naturally occurring metabolic changes that trigger enzymatic adaptation to maintain the function of the central nervous system [20]. Such metabolic changes are ideally subdued during electrophysiological experiments to establish tight control over neuronal signalling. In this work, the classical approach of mimicking the *in vivo* situation as closely as possible has thus been replaced by continuously improved attempts to suppress physiological side-effects from slicing and holding, such as excitotoxicity, trauma, hypoxia, edema, and nutrient deprivation, for as long as possible.

In the context of the 3R principle, the acute brain slice preparation can be considered an approach to *refine* and *replace* experiments that would otherwise be performed *in vivo*. Recent approaches have aimed to also *reduce* the number of animals used, by improving the viability of the preparation, thus allowing more and longer experiments [21–24]. Moreover, this allows to use more brain slices from a single animal by avoiding that intact brain slices have to be discarded at the end of a typical working day. For this purpose, a special incubation chamber ("Braincubator") was developed previously, in which pH and temperature of the perfusion medium are steadily controlled, and bacterial growth is suppressed by circulating the medium through a UV-C irradiated chamber [23]. While this obviously works, we found an alternative, simpler procedure by which neuronal viability is similarly increased.

Glucose was thought to constitute the only energy source for mammalian neurons [25] although it was shown that acute brain slices remained viable when glucose was replaced by lactate [26]. In the haemolymph of the leech, various organic anions, such as the Krebs-cycle components citrate, fumarate, lactate, and succinate, were found in mM concentrations [27,28], and changes in the composition of the haemolymph due to muscle activity and urine secretion have been described [29,30]. Whether these substances are utilized by neurons as energy sources remained unanswered. Instead, the functions of leech neurons and glial cells have always been investigated in “physiological” saline (“Ringer solution”), or standard leech saline solution (SLS) [3,4,6,7,31], often supplemented with glucose as the only organic substance. A single exception is one author’s diploma thesis [32], which reported that isolated segmental ganglia survive substantially longer in solutions containing one or more of the organic anions found in leech haemolymph.

Isolated leech neurons have also been cultivated [33,34], which provided insights into neurite outgrowth patterns, target recognition, and synapse formation [35]. For cultivation, Leibovitz-Medium L-15 was used [36,37] and under these conditions, the electrophysiological characteristics of neurons remained unchanged for ∼3 weeks [33,38]. However, the many advantages of the segmental ganglia, such as allowing to investigate neuron-glia-interactions [6], identifying functionally distinct neurons [4], or studying metabolic changes over time [3] are lost upon isolation and therefore, taking intact segmental ganglia into culture would be a desirable aim.

A key component of Leibowitz L-15 is pyruvate, which was also used in solutions for patch-clamp recording from neurons of adult mice [16]. When testing this preparation protocol, we found that in addition to being able to obtain recordings from neurons of adult mice, our slices additionally remained healthier for longer. We further optimized this protocol and introduced further procedures, until we found a method that enables to maintain viability for at least 36 hours. We show that the addition of pyruvate to the incubation media of acute brain slices allowed patch-clamp recordings on healthy neurons and coupled neuron pairs from mice aged P11 to ∼ two years for more than 36 hours after preparation, instead of the typical 8-12 hours. In contrast to similar advances using the Braincubator [21–24], our method builds on standard, albeit slightly modified incubation chambers and conventional technical equipment.

Given the potential benefit of our method, it should undergo independent assessment of other scientists in the field. We therefore wish to encourage further testing of our protocol to ensure its reliability.

## 2. Materials and Methods

### **a)** Segmental ganglia and identified neurons of the medicinal leech

#### Animals and preparation

The central nervous system of the leech consists of a head ganglion, a chain of 21 segmental ganglia, and a tail ganglion, which are connected by bundles of nerve fibres (“connectives”; see Fig. 1). A segmental ganglion contains ∼400 neurons [39]. The cell bodies are located in an outer layer (“outer capsule”) from which their extensions reach into the centre of the ganglion (“inner capsule”). There, they arborize to form synaptic contacts, thus constituting the so-called “neuropile”. From here, projections extend into the leech body via lateral roots, which are, again, bundles of nerve fibres over which signals are sent or received from the periphery. The outer capsule is divided into six compartments (“packets”), each of which contains a single packet glial cell, while the inner capsule contains two ‘giant’ neuropile glial cells. The segmental ganglia as well as connectives and lateral roots are enveloped by tough connective tissue which also separates the different compartments within the ganglia. The whole ganglion chain is located within a blood vessel (“ventral blood sinus”). Due to the stereotypical structure of the segmental ganglia, most neurons and glial cells can be identified based on their position in the ganglion, their size, and their electrophysiological properties.

Leeches were obtained from a commercial supplier (Biebertaler Blutegelzucht, Biebertal, Germany) and were identified as *Hirudo verbana* based on their dorsal and ventral coloration patterns [cf. 40]. They were kept in freshwater at 10 to 12 °C until used. Adult animals, measuring 3 to 5 cm in length when contracted and ∼9 to 13 cm when extended, were used for experiments, and segmental ganglia were prepared as previously described [3–5]. Briefly, leeches were outstretched and pinned down by their anterior and posterior suckers on a wax-covered surface. A deep incision behind the oral sucker severed the connectives between the first and second segmental ganglia, euthanizing the animal. The body was cut open along the dorsal midline and the central nervous system was dissected one segmental ganglion at a time (see Fig. 1), using fine scissors under a binocular stereomicroscope (Wild, Heerbrugg, Switzerland) [41]. Each segmental ganglion had ∼2 mm of connectives left attached to the anterior and posterior end of the capsule, and about 1 mm of the lateral root on either side (see Figs 1 & 3). The isolated segmental ganglia were transferred to covered glass dishes containing ∼2 ml of the respective incubation solution. These dishes were also stored overnight at 10 to 12 °C. To reduce microbial growth, glucose-containing medium was replaced daily. All other media were changed at least every three days.

### Incubation solutions

The choice of substances and composition of the incubation solutions (Table 1) used for maintaining leech ganglia is based on assessments of leech haemolymph [27]. Malate, which was not included in the original saline solution (abbr. BEL, from the German "Blutersatzlösung", which translates to ’blood replacement solution’) is in fact the dominant anion [28,30] with varying concentrations reported, depending on the state of the animal [27–31,42–44], feeding status, and stress [28–30]. Therefore, in preliminary experiments we tested four standard leech saline (SLS) solutions (see Table 1):

1. Standard leech saline (SLS), as used in our previous works [3–5]. Earlier works from Nicholls, Baylor, and Kuffler used "normal Ringer fluid" or "physiological saline" [31,45,46] that had an osmolality of ∼230 mOsm/kg, i.e. slightly hyperosmotic to leech blood. Later works used an SLS that was more isotonic to leech blood (∼200 mOsm/kg, [29,42]) and was similar to the one described here [3–5,47–49]. Some differences in ion composition, absolute concentrations, and pH buffering, had no major effect on experimental results.
2. SLS with 10 mM glucose (glucose-SLS) [cf. 7]. The various solutions used in previous works were also often complemented with glucose, but whether this affected neuronal function or the viability of the preparation has not been reported. Therefore, we here tested the effect of adding glucose to the SLS on the viability of the preparation (see below).
3. SLS with ∼15 mM malate (malate-SLS) as the only organic anion. Malate was found to be the most prominent organic anion in leech haemolymph, with concentrations ranging between 7 and 28 mM, depending on the state of the animal [27–31,42–44] and on feeding status and stress [28–30]. This solution had a markedly reduced [Cl^-^] (-30 mM), which was not compensated.
4. 5 mM pyruvate (pyruvate-SLS) as the only organic anion. The concentration of pyruvate was chosen from Leibovitz’s solutions [37], in which isolated neurons stay alive for weeks. It is noted that the concentrations of glucose and pyruvate in the leech haemolymph were found to be smaller than 1 mM [42,44]. Pyruvate was, however, repeatedly found to improve the preparation of brain slices [16,50] and it plays a major role in the "astrocyte-to-neuron lactate shuttle" (ANLS) hypothesis [51]. According to this, astrocyte-derived lactate is transported to neurons, where it is converted to pyruvate and used in the TCA-cycle as an energy source. Supplementing pyruvate via the holding solution renders this obsolete and may help keep lactate levels stable, while providing an unlimited supply of pyruvate (see Discussion). The slightly reduced [Cl^-^] (-5 mM) was not compensated.

**Table 1.**
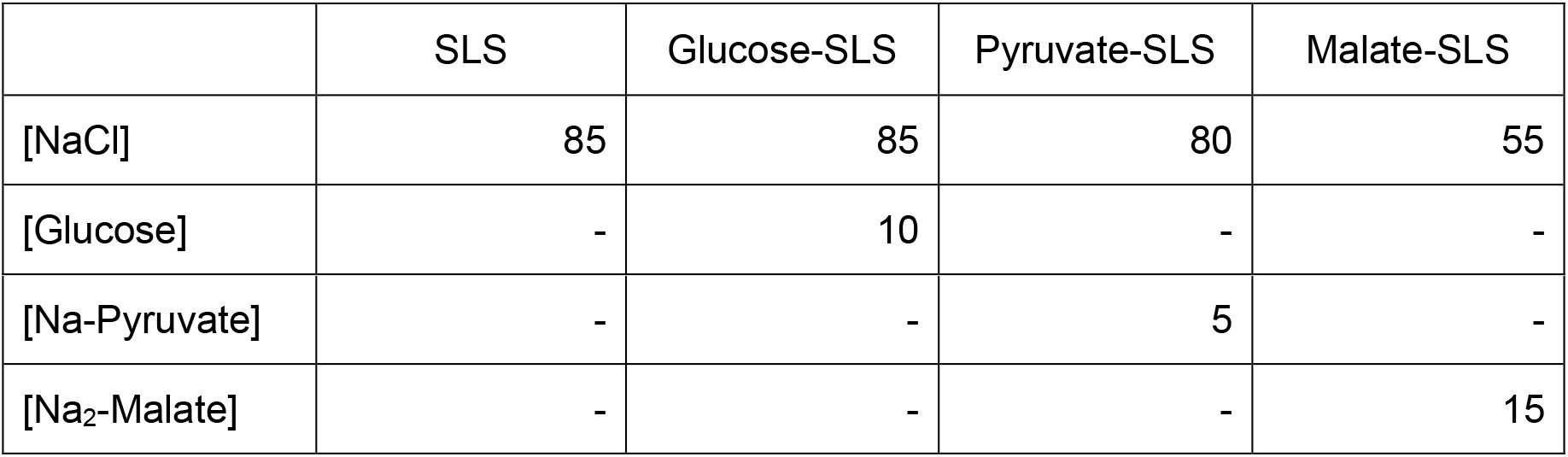
Composition (in mM) of the solutions used to prepare and incubate leech segmental ganglia. All solutions contained (in mM): KCl 4, CaCl_2_ 2, MgCl_2_ 1, HEPES 10; pH was adjusted to 7.40.

|  | SLS | Glucose-SLS | Pyruvate-SLS | Malate-SLS |
| --- | --- | --- | --- | --- |
| [NaCl] | 85 | 85 | 80 | 55 |
| [Glucose] | - | 10 | - | - |
| [Na-Pyruvate] | - | - | 5 | - |
| [Na<sub>2</sub>-Malate] | - | - | - | 15 |

These solutions were used for both the preparation and the subsequent incubation of isolated segmental ganglia. The glucose-SLS was passed through a sterile filter (Rotilabo, 0.22 μm pore diameter, Carl Roth, Karlsruhe, Germany) after preparation and stored in a fridge for a maximum of three days. All other solutions were stored in a fridge for a maximum of seven days. Solutions were prepared from stock solutions of NaCl, KCl, CaCl_2_, MgCl_2_, HEPES (N-(2-Hydroxyethyl)piperazine-N′-(2-ethanesulfonic acid)), Na-pyruvate, and Na_2_-malate. Chemicals were from VWR (Leuven, Belgium; NaCl, KCl), Acros Organics (Geel, Belgium; CaCl_2_), Invitrogen (Carlsbad, CA, USA; Na-pyruvate), Sigma-Aldrich (St. Louis, MO, USA or Seelze, Germany; 5-Hydroxytryptamine, KOH), Grüssing (Filsum, Germany; MgCl_2_, KCl), Merck (Darmstadt, Germany; NaOH), AppliChem (Darmstadt, Germany; HEPES, Na_2_-Malate), Carl Roth (Karlsruhe, Germany; HEPES), or Caelo (Hilden, Germany; Glucose-Monohydrate). The pH was set to 7.4 by using NaOH. The osmolality of the solution was verified by a freezing point depression osmometer (Fiske Model 210, Advanced Instruments Inc., Norwood, MA, USA or Osmomat 030, Gonotec, Berlin, Germany).

We tested the viability of segmental ganglia by assessing the transparency of the ganglion capsule, spontaneous action potential generation and resting E_m_ in so-called Retzius and Leydig neurons, as well as E_m_ of the neuropile glia cells (Fig. 2-3). To test for the effects of reduced [Cl^-^]_o_, serotonin (5-Hydroxytryptamine, 5-HT, 1 mM) was applied in a subset of experiments via the bath. 5-HT increases Cl^-^conductance in leech Retzius neurons [52] and therefore induces E_m_ changes according to E_Cl_.

**Fig. 2:**
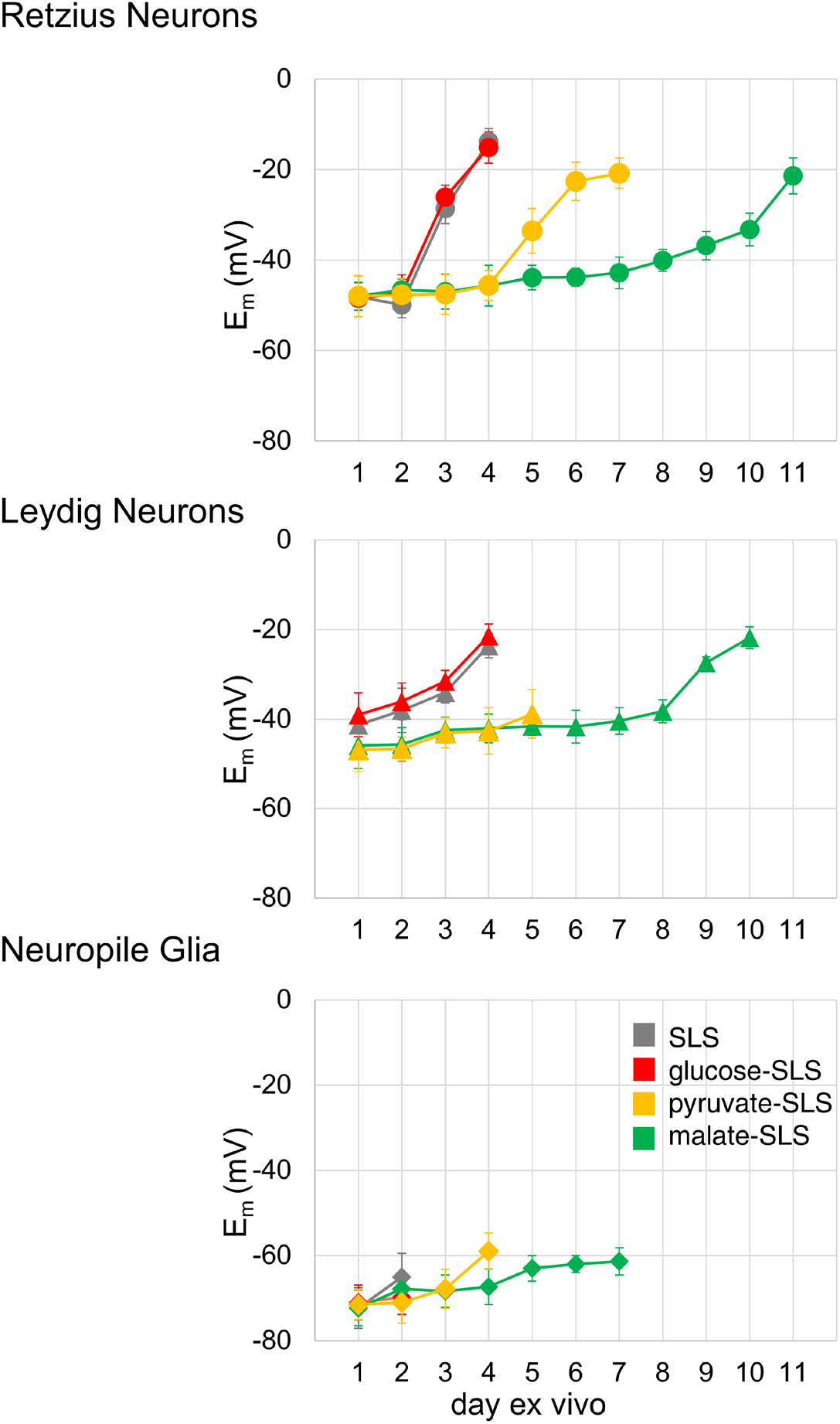
E_m_ of leech neurons and neuropile glia cells. Resting E_m_ of Retzius neurons, Leydig neurons, or neuropile glia cells, as measured at different days after the preparation at day 1 (’days ex vivo’). The ganglia were prepared, stored, and used for recordings either in standard leech saline (SLS) or in SLS complemented with 10 mM glucose, 5 mM pyruvate, or 15 mM malate. Data points are mean ± SD of n = 4 to 13 neurons and of 3 to 5 glial cells.

### Electrophysiological recordings of identified leech neurons and glia cells

Glass microelectrodes were pulled from borosilicate glass capillaries (OD 1.5 mm, ID 0.86 mm, length 75 mm, Harvard Apparatus, Edenbridge, UK) on a vertical puller (PE-2, Narishige, Tokio, Japan) and filled with 0.5 M K_2_SO_4_ (JT Baker, Deventer, The Netherlands) and 20 mM KCl (series resistance ∼60 MΩ). A chlorided silver wire (dia. 0.5 mm) was inserted into the electrode and the assembly was sealed with hard wax (Deiberit 502, Böhme & Schöps Dental, Goslar, Germany). Electrodes were connected to a custom-made amplifier (Central workshop Biology, Waldemar Seidel, Heinrich Heine University Düsseldorf), which in turn was connected to an AD converter (Digidata 1322A, Molecular Devices, Axon Instruments, San Jose, CA, USA). The experimental chamber was grounded by an Agar bridge made from plastic tubing (OD 3 mm, ID 2 mm, length 40 mm) filled with 3% Agar in 3 M KCl that had a chlorided silver wire inserted into it and which was positioned close to the chamber outflow to avoid contamination of the superfusion medium with KCl. The silver wire was not in direct contact with the bath solution.

Segmental ganglia were transferred into an acrylic chamber, filled with a silicone elastomer (Sylgard 184, Dow Corning, Midland, MI, USA) to which the preparation was pinned. The bath volume was ∼130 μl and perfusion was gravity driven at a rate of ∼14 μl/s, exchanging the bath every ∼9 s.

Electrode placement was performed using a micromanipulator (Type 115, Leitz, Wetzlar, Germany) under optical control with a stereo microscope (Helmut Hund GmbH, Wetzlar, Germany). The entire setup was placed on a vibration-dampened steel plate inside a grounded Faraday cage.

Ganglia were fixed ventral side up and leech neurons were identified by their position and by their electrophysiological characteristics. Typically, after inserting the electrode, spontaneous action potentials were observed and their frequency decreased over the following few minutes before reaching a relatively stable value. Resting E_m_ was determined subsequently after it had stabilized within a range of ±1 mV. Neuropile glia cells were accessed by inserting the electrode in between the two Retzius neurons deep into the ganglion and identified by their comparably negative resting E_m_ or by their marked depolarizing response to 40 mM K^+^ in the superfusion medium. All experiments were conducted at room temperature. Experimental data were analysed every day for up to eleven days after preparation in the four incubation solutions (Table 1). Experiments continued every day until we could no longer obtain stable recordings.

### **b)** Experiments on neurons of acute brain slice preparations of mice

#### Animals and preparation

All animal procedures were in accordance with the guidelines of the European Community’s Council Directive of 22 September 2010 (2010/63/EU). Animal euthanasia for brain removal was registered with the regional council (Regierungspräsidium Freiburg, X-17-05C and X-22-09B) and all animal procedures were approved by the Center for Experimental Models and Transgenic Service (CEMT, University Freiburg) and by the local authorities (Regierungspräsidium Freiburg, G-19/07).

Mice (*Mus musculus*, C57BL/6J, from Janvier or Charles River) were housed in individually ventilated and air-conditioned cages under a 12 h light-dark cycle (light off 10:00 -22:00), in a temperature- and humidity-controlled room. Litters were housed together with a dam until weaning and separated at postnatal day (P) 21 to 23. Food and water were provided *ad libitum*.

Experiments were performed on neurons from

I. layer 1 (L1) of the medial prefrontal cortex (mPFC, separated into mice aged P11–32 and P33+),
II. L1 of the motor cortex (M1/M2, P15-32),
III. globus pallidus (GP, P38+), and
IV. thalamic reticular nucleus (TRN, P11-36).

In some cases, slices and neurons from the dams were included, which were ∼1-2 years of age at the time (see results).

Mice were anaesthetized with isoflurane (2% induction, gradual increase to 5%, with 95% O_2_) until loss of tail-flick and foot-withdrawal reflex. For animals aged P33 or older, this was followed by a cervical dislocation that left the carotid arteries intact and by a transcardial perfusion with recovery-ACSF. For younger mice, aged P11 to P32, anaesthesia was followed directly by decapitation with a small-rodent-guillotine (Type 7575, Hugo Sachs Elektronik - Harvard Apparatus GmbH, March, Germany). We found this method (four-blade iris cut) to be superior to using scissors: Due to symmetric cutting and fixation, brain trauma by one-sided pressure is prevented, in particular for animals older than ∼P18.

Following decapitation, the head was placed on a frozen metal platform (previously taken from its storage at -21 °C) and doused in ice-cold recovery-ACSF containing (in mM): NMDG, 92; HCl, 93; KCl, 2.5; NaH_2_PO_4_, 1.25; NaHCO_3_, 30; HEPES, 20; glucose, 25; L-ascorbic acid, 5; thiourea, 2; sodium pyruvate, 3; MgSO_4_ • 7H_2_O, 10; N-acetylcysteine, 12; CaCl_2_, 0.5; pH 7.35 was adjusted with HCl, osmolality was adjusted to ∼305 mOsm/kg, and the solution was gassed with carbogen (95% O_2_, 5% CO_2_).

A block of brain containing either GP and TRN and/or mPFC and M1/M2 was promptly removed after decapitation and submerged in ice-cold recovery-ACSF (see Table 2). Coronal slices of 350 μm (for animals aged P11-32) or 400 μm thickness (for animals aged P33 and older) were prepared in this solution with a vibratome (VT1000S microtome, Leica, Germany). During this step, gassing with carbogen was commenced in between the slicing. Acute brain slices were then carefully collected with a fire polished pipette (∼4 mm inner diameter) and transferred into one of two types of holding chambers (see Fig. 4A):

1. BSK-12-chamber (Brain slice keeper 12, Science Products GmbH, Hofheim, Germany): This holds 500 ml and contains 12 wells of 0.3 cm depth and 1.2 cm diameter each.
2. UKM-chamber (our own holding chamber optimized to improve the viability of the slices, see acknowledgements): This holds 550-600 ml and has 8 wells of 1.6 cm depth and 1.8 cm diameter each).

**Table 2.** Composition (in mM) of the solutions used to prepare, incubate, and record from mouse acute brain slices. pH was adjusted to 7.35 using HCl for recovery-ACSF or NaOH for the other solutions. Osmolality was adjusted to ∼305 mOsm/kg.

|  | recovery-ACSF<br>(slicing) | pyruvate-ACSF<br>(holding,<br>incubation) | standard-ACSF<br>(recording) | comment |
| --- | --- | --- | --- | --- |
| NaCl | - | 85 | 124 |  |
| NMDG-Cl | 92 | - | - | impermeable Na<sup>+</sup> replacement; shuts down neuronal activity; prevents hypoxic conditions and ATP depletion during slicing [50,53]. Produced by titrating NMDG with HCl. |
| KCl | 2.5 | 2.5 | 2.5 |  |
| MgSO<sub>4</sub> | 10 | 2 | 2 | higher concentration of Mg<sup>2+</sup> shuts down neuronal activity by NMDA receptor block [18,55]. Used as a heptahydrate. |
| CaCl<sub>2</sub> | 0.5 | 2 | 2 | lower concentration reduces Ca<sup>2+</sup> second messenger signalling [55] |
| NaH<sub>2</sub>PO<sub>4</sub> | 1.25 | 1.2 | 1.2 | weak buffer |
| NaHCO<sub>3</sub> | 30 | 26 | 24 | maintains pH in combination with carbogen |
| HEPES | 20 | 20 | 5 | CO<sub>2</sub>-independent buffer, reduces edema [16] |
| Glucose | 25 | 20 | 12.5 | usefulness unclear [25,26] |
| L-ascorbic acid | 5 | 5 | - | counters oxidative stress during slicing and incubation [56,57], reduces edema [16,57], possibly antimicrobial effect |
| Thioruea | 2 | 2 | - | complexation of metals, reduces edema [16], possibly antimicrobial effect, antioxidant in conjunction with ascorbate [56] |
| Na-pyruvate | 3 | 3 | - | maintains energy metabolism, prevents excitotoxicity [54] |
| N-acetylcysteine | 12 | 12 | - | counters oxidative stress during slicing and incubation [16], possibly antimicrobial effect |

Deeper wells are an advantage when moving the incubation chamber and help to keep slices contained therein in position. Moreover, deeper wells should help to enable linear flow of gassed solutions and the larger volume per slice may be beneficial for overnight storage (see below) by reducing the effects of accumulated metabolic end products. Both chambers ensured continued gassing with carbogen and secure superfusion of slices during the holding/incubation procedure.

### Solutions and incubation

The chambers were completely filled with pyruvate-ACSF. The original composition of pyruvate-ACSF contains (in mM): NaCl, 92; KCl, 2.5; NaH_2_PO_4_, 1.2; NaHCO_3_, 30; HEPES, 20; glucose, 25; L-ascorbic acid, 5; thiourea, 2; sodium pyruvate, 3; MgSO_4_ • 7H_2_O, 2; N-acetylcysteine, 12; CaCl_2_, 2; pH 7.35 was adjusted with HCl, osmolality was adjusted (by adding H_2_O) to 300 to 305 mOsm/kg, and solution was gassed with carbogen. This pyruvate-ACSF underwent refinement in NaCl, NaHCO_3_ ^-^, and glucose content in order to reduce or avoid osmolality adjustments (modified pyruvate-ACSF), and contained: NaCl, 85; KCl, 2.5; NaH_2_PO_4_, 1.2; NaHCO_3_, 26; HEPES, 20; glucose, 20; L-ascorbic acid, 5; thiourea, 2; sodium pyruvate, 3; MgSO_4_ • 7H_2_O, 2; N-acetylcysteine, 12; CaCl_2_, 2; pH 7.35 was adjusted with HCl, osmolality required very little addition of water to reach 300-305 mOsm/kg. The solution was gassed with carbogen.

The pyruvate-ACSF in the chambers was heated to 32 °C prior to the slicing procedure. For this, the BSK-12-chamber was placed in a heated water bath (PolyScience WB05, Niles, IL, USA), and the UKM-chamber was placed into its base (Fig. 4C, black), where it remained for the rest of the day. Slices were incubated at 32 °C by the heated base or the water bath for 20-40 min, and were then allowed to equilibrate at 22 °C for at least 60 min before transferring them, one at a time, into the recording chamber. For cooling down to room temperature, the BSK-12-chamber had to be removed from its water bath. For the UKM-chamber, the power to the heating element was simply turned off but the chamber remained stationary.

We hypothesized that the modified pyruvate-ACSF prolongs the viability of slices and allows experiments on brain slices after overnight storage. Thus, slices not used on the day of preparation were stored overnight in pyruvate-ACSF. The holding chamber was carefully transferred into a fridge. This fridge was adjusted such that ice crystals just formed on the evaporator but temperature at the position of the holding chamber was entirely above freezing and as close as possible to +2 °C. This was monitored with thermometers inside the compartment next to the chamber position. Although gassing was discontinued inside the fridge, the chamber should be saturated with carbogen and, with the lid on, the pH of the pyruvate-ACSF was mostly stable until the next day, when gassing commenced. The large volume of the holding chamber, the space above the solution (filled with carbogen), the lid (limiting gas exchange), and the cooling could prevent a lack of O_2_ and emission of CO_2_. On the next morning, the holding chamber was carefully removed from the fridge, immediately gassed again with carbogen, and then allowed to equilibrate to room temperature (∼22 °C, ∼1 h). Slices were used throughout the entire day (day 2) until 36 hours after preparation. Some slices were also used on day 3 or day 4, as indicated.

All solutions described here were prepared with deionized water with a specific resistance of 18.2 MΩ•cm and filtered through bottle-top filters into sterile bottles (Steritop vacuum filtration system, 0.22 μm pore size, Millipore, Merck KGaA, Darmstadt, Germany).

Data collected from the TRN and the mPFC are from slices incubated in both holding chambers and original and modified pyruvate-ACSF, but are pooled in analysis. However, using the UKM-chamber and the modified pyruvate-ACSF improved our subjective success rate for obtaining results from viable slices and healthy neurons on day 2 after preparation. For example, when moved to cold storage, the UKM-chamber did not cause slices to swirl around and swim first to the surface and then to the bottom of the chamber, as was frequently observed using the BSK-12-chamber. We considered the lack of this very annoying effect an enormous improvement. Since we did not monitor or analyse our unsuccessful attempts or ruined incubations, we have no data supporting the claim of an improved success rate and therefore this statement should be regarded as merely anecdotal.

### Presumed function of individual components in the solutions

The recovery-ACSF, based on NMDG (see Table 2), is used to protect neurons during acute brain slice preparation and was inspired by previous work (Jonathan T. Ting, personal communication and [16]). Since Na^+^ channel pores are not permeable to methylated organic cations [53], such as NMDG^+^, neuronal signalling is shut down, and thus, hypoxic conditions are counteracted during the preparation. Slicing in ice-cold NMDG solution was found to be necessary for maintaining neuronal viability in several cell types, brain regions, and animal ages. It is based on the assumption that a chain of events is started by Na^+^ influx: depolarisation → Cl^-^ influx → water entry → cell swelling. It was suggested that this is the predominant reason for poor neuronal survival during slice preparation [16,50]. Additionally, L-ascorbic acid and N-acetylcysteine are added to both the recovery-ACSF and the pyruvate-ACSF to counter oxidative stress due to depletion of cellular antioxidants during the slicing and incubation procedures [16]. Supplementing substances that help to maintain cellular energy metabolism can additionally prevent delayed excitotoxicity [54].

Exogenously applied pyruvate was shown to restore ATP levels and prevent delayed neuronal degeneration and synaptic deterioration when administered after NMDA receptor activation [54]. NMDA receptor activation during the rewarming phase can be counteracted by omitting Ca^2+^ from solutions and using instead high concentrations of Mg^2+^ (10 mM) as described previously [18,55]. The final solutions are summarized in Table 2.

### Electrophysiological recordings

For whole-cell patch clamp recordings, a continuous single-electrode amplifier was used for both current injection and E_m_ measurement (Axon Instruments MultiClamp 700B driven by a desktop computer through a Digidata 1550B AD converter, Molecular Devices, San Jose, CA, USA). All experiments were performed in an air-conditioned room at 22 °C. The experimental setup was housed in a Faraday cage for shielding against electrical interference. The entire setup was mounted on an optical table with tuned vibrational isolators (Newport RS4000, Newport, Irvine, CA, USA) for dampening mechanical vibrations. A single slice was transferred into the recording bath chamber (Slice Mini Bath Chamber I, Luigs & Neumann, Ratingen, Germany) and secured in place with a U-shaped slice anchor with nylon cross strings (Slice Anchor - Flat Stainless Steel, 2.0 mm, SHD-40/2, Multichannel Systems GmbH, Reutlingen, Germany). The recording bath chamber was filled with standard-ACSF solution containing (in mM): NaCl, 124; KCl, 2.5; NaH_2_PO_4_, 1.2; NaHCO_3_, 24; HEPES, 5; glucose, 12.5; MgSO_4_ • 7H_2_O, 2; CaCl_2_, 2; pH 7.35 was adjusted with HCl, osmolality was adjusted to 300-305 mOsm/kg, and solution was gassed with carbogen. This solution was also used as standard-ACSF when comparing incubations between pyruvate-ACSF and standard-ACSF (see below).

Rate of flow of standard-ACSF into the chamber was ∼4-6 ml/min. The bath chamber was grounded via a chlorinated silver wire or an Ag/AgCl-pellet submerged in the standard-ACSF. Glass pipettes were pulled from borosilicate glass capillaries with filament (BF150-86-10, Sutter Instrument, Science Products) using a vertical pipette puller (PIP-6, HEKA Elektronik, Multichannel Systems GmbH) and were filled with electrolyte solution containing (in mM): NaCl, 10; K-gluconate, 105; K_3_-citrate, 20; KCl, 1; HEPES, 10; EGTA, 0.5; Mg-

ATP, 3; Tris-GTP, 0.5; MgCl_2_, 0.5; pH was adjusted to 7.30 with KOH, and osmolality was adjusted to 295-300 mOsm/kg. We found that results were improved and recordings more stable over time when the osmolality of the electrolyte was slightly lower than that of the bath solution. Hypothetically, a higher intracellular osmolality would lead to the inflow of water to equilibrate the osmolality. However, since the electrolyte volume in the patch pipette (∼20 μl) is larger by five to six orders of magnitude than the intracellular volume (5-100 pl), this is a futile mechanism when using low resistance electrodes in the whole-cell configuration. Dilution by water inflow would be counteracted by the electrolyte. Extruding ions in regulatory volume decrease [3] is again futile, since the entire electrolyte would have to be cleared of excess osmolytes. Shrinkage, however, is usually self-limited due to non-mobile osmolytes: a volume decrease by an outflow of water causes a slight concentration increase.

The Ag/AgCl wire was submerged into the filled glass pipette, completing the microelectrode assembly. Electrode resistances after submersion in the bath were between 3.5 and 8 MΩ. Positioning of these recording electrodes was performed using micromanipulators (LN Junior Unit with SM10-Remote Control System, Luigs & Neumann, Germany).

Location of the area of interest in the brain slice was estimated using the Interactive Mouse Brain Atlas (Allen Institute, Seattle, WA, USA) and a 5x objective lens (Olympus MPlanFL N 5x/0.15 NA). Neurons were identified and accessed under optical control using a 40x or 20x objective lens (Olympus LUMPlanFLN 40x/0.80 NA or XLUMPlanFLN 20x/1.0 NA). Recording electrodes were offset to zero mV in the bath solution before each experiment. Whole-cell patch clamp configuration was achieved in voltage-clamp mode with MultiClamp Commander (version 10.7, Axon Instruments Inc.). Capacitance was compensated (slow and fast pipette capacitance compensation, MultiClamp 700B). We additionally attempted to minimize remaining transients by reducing the capacitance from the immersed and filled surface area of the micropipette (i.e. reducing all fluid levels), by reducing stray capacitance from the remaining surface using glass capillaries with the thickest wall possible, and by using an electrode holder optimized for low stray capacitance (Luigs & Neumann, Ratingen, Germany). Afterwards, the remaining capacitances and series resistances were compensated and the values noted as C_in_ (see results) and R_s_. All following recordings were made in the current clamp mode with adjusted bridge balance. The membrane potential of each cell was adjusted to -60 or -70 mV by an additional DC current offset. Injections of DC current (I_cc_) were used to obtain I/V relations, which were used to further analyse input resistance (R_in_). Changes of R_in_ over time were used to monitor activation or inactivation of ion channels. Currents were injected for a duration of 500 ms (current step) before returning to the Em of -60 or -70 mV (with manual DC offset as adjusted previously) for 1.5 to 2.5 s (sweep). Current steps were chosen such that they changed the membrane potential of neurons in step-wise incremental increases. Thus, the current injections changed from hyperpolarizing to depolarizing and the membrane potential changed from ∼-100 mV in the first sweep to a depolarized membrane potential that produced continuous action potential generation in the last sweep. The frequency of the APs greatly depended on the injected current and when depolarization blocks were observed, the recording of the I/V relation was considered complete. We found that injecting more depolarized currents beyond the point of depolarization blocks may damage neurons.

The liquid junction potential was not corrected for (calculated to V_m_ = V_P_ - 14.2 mV; considering all ions as free; with V_m_ being the true membrane potential without "zeroing" offset, and V_p_ being the recorded membrane potential; calculated using LJPcalc [58] and a stationary Nernst-Planck equation).

### Imaging of Cell Viability in pyruvate-ACSF compared to standard-ACSF

To assess cell viability, freshly prepared brain slices were incubated and stored during the day, either in standard-ACSF (slices were from an animal aged P23) or in pyruvate-ACSF (P17). A single brain slice was then removed from the incubation chamber and transferred to a small volume staining chamber (5 ml) where it was incubated for 20 min at RT in fresh carbonated standard-ACSF supplemented with 1 ml solution of the LIVE/DEAD Cell Imaging Kit (two colour kit with excitation at: 488 and 570 nm and emission at 512 (green) and 620 nm (red); Invitrogen/ThermoFisher Scientific, Darmstadt, Germany). The 5 ml staining solution was freshly prepared for every slice. This kit contains the cell permeant dye Calcein-AM, as an indicator for living cells (requiring intact esterase activity), and the nuclear staining dye BOBO-3 iodide, as an indicator for dead cells (with a compromised membrane). After incubation with the respective dye, the single slice was removed from the small volume staining chamber and placed into the recording chamber of the microscope. Here, it was superfused with carbogenated standard-ACSF for 15 min, to wash out any remaining dye.

Staining and subsequent imaging was performed either on the day of preparation (day 1), or 24 h (day 2), 48 h (day 3), or 72 h (day 4) afterwards. All slices were from the same original preparation, but slices were freshly incubated in the dyes immediately before imaging. We continued to stain and image slices from the incubation chamber until no more living (green) cells were found. This occurred on day 3 for the slices incubated in standard-ACSF, and on day 4 for the slices incubated in pyruvate-ACSF. Cells were counted and analyzed in L2/3 of the mPFC. For comparison with the electrophysiological assessment of cell viability, L1, TRN, or GP neurons would have been preferable, but we found either the cell density or tissue contrast to be too low for a meaningful comparison of incubation solutions.

For each image, four channels were recorded: upper green channel (UG: bandpass filter 525/50 nm, Brightline HC, AHF Analysentechnik AG, Tübingen, Germany), upper red channel (UR, bandpass 641/75 nm, Brightline HC, AHF), lower green channel (LG, 490-550 nm bandpass filter, Femtonics, Budapest, Hungary) and lower red channel (LR 575-650 nm bandpass filter, Femtonics). A two-photon laser (Chameleon Discovery, Coherent Corp., Saxonburg, PA, USA) tuned to 800 nm was used, which excited both fluorophores. A λ/2 plate and a Glan Thompson Polarizer were used for pre-attenuation of the laser, such that the pockels cell, which followed in the optical path, could operate at medium voltages, where spot quality was found to be best. The Pockels cell (Conoptics, Danbury, CT, USA) was used for fast final attenuation of the laser. Green GaAsP PMTs were set to 70% Gain, red multialkaline PMTs to 60%, to obtain an optimal signal-to-noise ratio at lowest possible laser powers. In the experiments, image brightness was adjusted by using the motorized λ/2 plate mount or by adjusting the voltage to the pockels cell driver.

### Data Analysis

#### Imaging

Image analysis to confirm cell viability was performed using ImageJ (Fiji bundle, licensed under the GPL). For every time point (Days 1-4), six to ten areas of interest of 90 x 90 μm and a depth (along the z-axis) of 30 μm were analysed. For area analysis, the channels were split and a 50-pixel rolling-ball was used to subtract the background for each channel (ImageJ). Then, the lower and upper channels were summed (UG+LG, UR+LR). A one-to-two-pixel Gaussian blur was applied to the resulting images. For subsequent ROI detection, a threshold was used to generate the ROI masks. This was set at the point of the histogram beyond which a rapid decay to zero occurred, i.e. at the point of transition from the main peak to the tail of the distribution. When cells were positioned very close to one another, the threshold was adjusted to maximize coverage of each cell, but without causing masks to merge. If any masks still fused despite this adjustment, a watershed algorithm was applied to separate the objects. Cells were counted based on the resulting masks, where objects with an area of less than 10 μm^2^ were removed.

The number of cells was counted for the green (alive, Calcein-AM) and red (dead, BOBO-3 iodide) channels separately. Any cluster of small objects (thus not clearly a cell body) marked by Calcein-AM, but adjacent to a red-stained nucleus, was still considered a dead cell, as it obviously represents Calcein-encasing membrane debris (green) of a former cell with a now free nucleus (red). The percentage of alive cells among the total number of cells was calculated (Fig. 6A). Additionally, the size of the ROIs was calculated in order to monitor cell bloating or swelling and to thus assess a potential precursor to cell death (Fig. 6B). Finally, the number of cells per area of 100 × 100 μm (calculated from the 90 x 90 μm images, Fig. 6C) was calculated to be able to compare cell density.

Within ACSF groups, differences were statistically assessed using the Kruskal-Wallis test followed by Dunn’s post hoc test with Holm correction for multiple comparisons. Between ACSF groups, pairwise differences were analysed using the Mann-Whitney test with Holm-adjusted p-values.

#### Electrophysiology

For analysing electrophysiological data, Clampfit 10.7 software (Axon Instruments, Sunnyvale, CA, USA) and Python (Python Software Foundation, Wilmington, DE, USA) with pyABF package (https://swharden.com/pyabf/) and our own customized scripts were used: [https://github.com/IMBIT-Cellular-Neurophysiology-Freiburg/ElectrophysData]. Graphical illustrations of the results were done in OriginPro (version 2021, OriginLab Corp., Northampton, MA, USA), and Excel (version 2016, Microsoft Corp., Redmond, WA, USA).

Cells were included in the analysis when they had a resting E_m_ more negative than -48 mV, an access resistance R_a_ in voltage-clamp mode smaller than 25 MΩ, and a bridge balance in the current-clamp mode smaller than 20 MΩ. All data were subdivided into groups of the day since slicing on which the experiment was conducted, i.e. the day of preparation was day 1, after one overnight incubation it was day 2, and, in some cases, we also obtained data from day 3. All results are expressed as mean ± SEM. Resting membrane potential (E_m_) was recorded in current-clamp mode when no DC current injection was applied. Input resistance (R_in_) was calculated from the slope of a part of the I/V relation that rendered E_m_ between -80 mV to -50 mV. To determine C_m_ a small current injection was used that hyperpolarized E_m_ by at least 10 mV to ∼-80 mV, avoiding any impact of I_h_ from HCN channels. The time constant τ (tau) was determined when 1/e of the total voltage change induced by this current step had occurred, i.e. the time passed from the beginning of the current step until a voltage of E_τ_ = ((E_m_ - E_cc_)/e) + E_cc_) was reached. The membrane capacitance C_m_ was calculated from this as C_m_ = τ/R_in_. Additionally, the input capacitance (C_in_) was determined at the beginning of each experiment, when determining whole cell access resistance under voltage clamp conditions. The values were considerably different between these two techniques. Reasons for this may include but are not limited to the multidimensional nature of the capacitance of dendritic compartments, hyperpolarization-activated ion channels that have activated already at -80 mV, or inactivating ion channels that were active at rest and have inactivated at -80 mV.

Action potential firing frequencies were calculated near the threshold, when the depolarizing current injection was sufficient to cause at least 4 action potentials. The initial frequency (i.e. between the first two spikes), late (between last two spikes) and mean frequency (between all but the first two spikes) were calculated from peak-to-peak durations. For cells in the TRN, we additionally determined the width of low-threshold spikes (LTS), which are depolarizing Ca^2+^-envelopes of ∼200 ms duration, typically crowned by action potentials [59–61]. We analysed them following release from a hyperpolarizing injection step. We determined their duration by using the first and last inflection point on either side of the LTS event. The firing frequency was calculated for all spikes in an LTS.

Normal distribution was assessed using the Shapiro-Wilk test. When the data were not normally distributed, comparisons were performed using a two-sided Mann-Whitney U test. When data were normally distributed, a student’s t-test was used. For two-group comparisons the p-values for significance were p < 0.05 (*), p < 0.01 (**), and p < 0.001 (***). For comparisons involving three groups, pairwise Mann-Whitney U tests were applied with Bonferroni-adjusted p-values. For categories (e.g. LTS propensity 0/1) we used Fisher’s exact test.

## 3. Results

### **a)** Effect of malate and pyruvate on leech ganglia, neurons, and glia cells

Since the central nervous system of the leech is embedded in the ventral blood sinus (see Fig. 1A), it is feasible to assume that energy metabolism is sustained by one or several haemolymph components. We therefore assessed how long leech segmental ganglia are viable in our previously used SLS [3,4,6,7,31], as well as in glucose-SLS, pyruvate-SLS, or malate-SLS (see Table 1). Spontaneous AP generation (neurons) after first electrode insertion and E_m_ (neurons and neuropile glia cells) were analysed every day for up to eleven days after preparation in the four incubation solutions.

When stored overnight at room temperature and investigated the following day (day 2), Retzius neurons no longer generated spontaneous APs and were strongly depolarized (not shown). When stored at 10-12 °C overnight, the neurons also failed to generate spontaneous APs on day 2, but were not depolarized until day 3 (Fig. 2, top panel). The same observation was made using glucose-SLS, which shows that glucose is ineffective in sustaining the viability of isolated leech ganglia. In contrast, in solutions containing 5 mM pyruvate, APs in Retzius neurons were still generated on day 3, and in 15 mM malate, APs were generated spontaneously until day 8. Typically, Retzius neurons spontaneously generated ∼4 APs in 10 s with amplitudes of ∼20 mV. In Leydig neurons, APs were generated until day 3 in all solutions except malate-SLS, where spontaneous AP generation continued until day 8. Typically, Leydig neurons spontaneously generated 1-7 APs in 10 s with amplitudes between 30 and 35 mV. Frequency and amplitude of spontaneous APs in Retzius and Leydig neurons varied considerably between ganglia and between cells of the same ganglion. We could not attribute any changes in AP frequency or time course to the number of days *ex vivo* or on the incubation solution.

Retzius neurons gradually depolarized over time, initially with a slow rate of less than 1 mV per day and then considerably faster after day 2 in SLS and glucose-SLS, after day 4 in pyruvate-SLS and after day 7 in malate-SLS. In Leydig neurons, E_m_ was depolarized in SLS and glucose-SLS from the start, while it gradually depolarized in the other solutions. An E_m_ negative to -40 mV was maintained by Leydig neurons in pyruvate-SLS until day 5 and in malate-SLS until day 9 (Fig. 2, middle panel).

We were able to record E_m_ from neuropile glia cells in SLS and in glucose-SLS on the day of preparation and the day after (Fig. 2, bottom panel). In pyruvate-SLS, we were able to record up to day 4. In malate-SLS, it was up to day 7. E_m_ of neuropile glia cells gradually depolarized over time. In contrast to neurons, electrophysiological access to the neuropile glia cells is severely hampered by the state of the ganglion capsule (see Fig. 3). Therefore, the inability to access neuropile glia cells is not necessarily an indication of its state of viability. In summary, leech segmental ganglia that had been stored refrigerated and in pyruvate- or malate-SLS could be used at least over three days without obvious constraints to E_m_ or spontaneous AP activity.

**Fig. 3:**
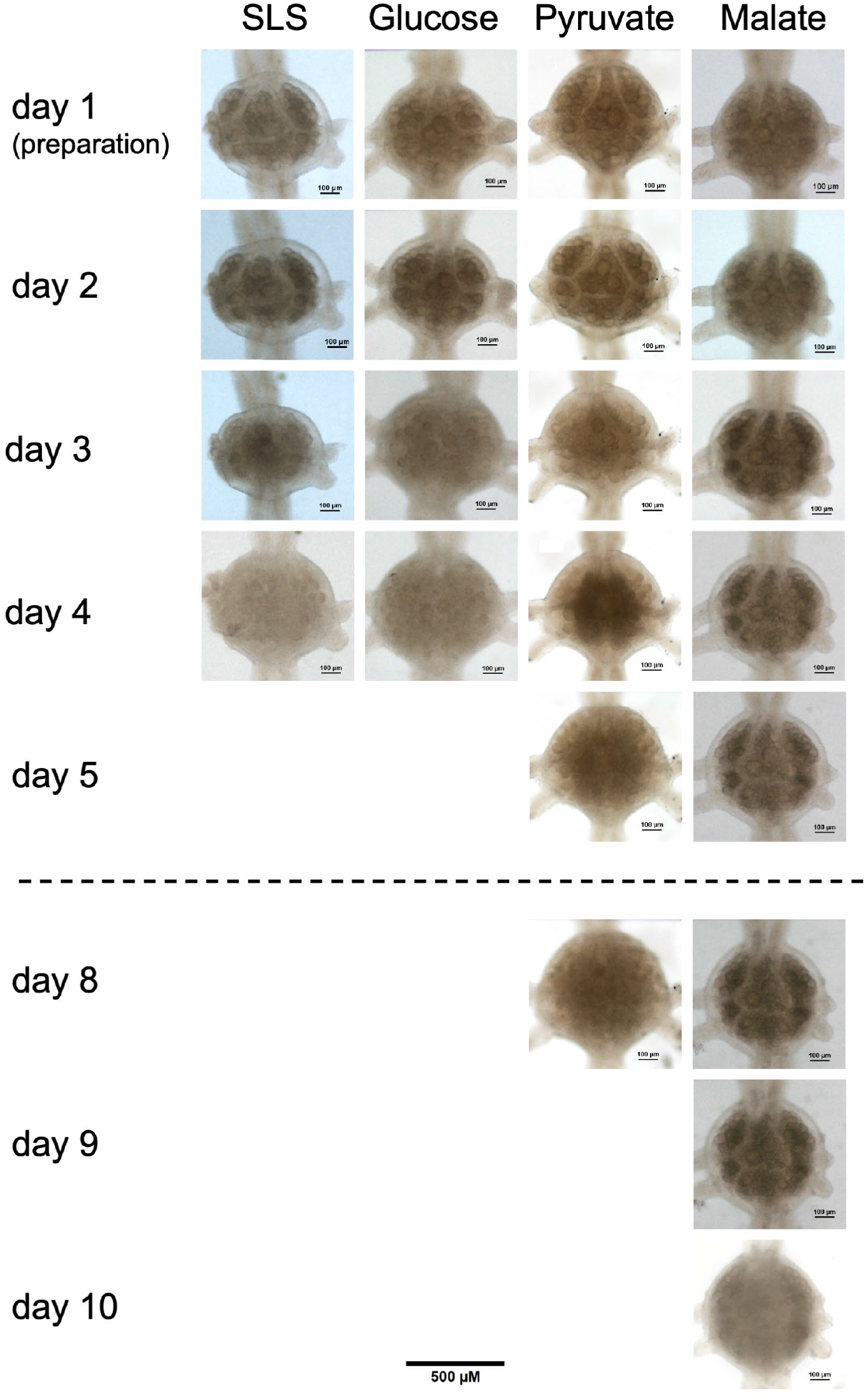
Effect of *ex vivo* conditions on the ganglion capsule. The transparency of the ganglion capsule was also visibly inspected to assess viability. In SLS and glucose-SLS, the capsule became opaque on day 3, in pyruvate-SLS on day 5, and in malate-SLS on day 10. Figure scale bar is 500 μm, the scale bars in the images are 100 μm.

In malate-SLS, the Cl^-^ concentration was considerably lower than in the other solutions, which should affect the Cl^-^ equilibrium potential (E_Cl_) and therefore, E_m_. However, exchanging the bath perfusion medium from SLS to malate-SLS for up to 3 min did not change the resting E_m_ or AP generation in Retzius or Leydig neurons, suggesting a very low Cl^-^ permeability. We therefore tested the application of serotonin (5-Hydroxytryptamine, 5-HT), which increases Cl^-^ conductance in leech Retzius neurons [52], and thereby induces E_m_ changes proportionate to E_Cl_. Exposure to 1 mM 5-HT caused hyperpolarizations in SLS to -56 ± 3 mV (n = 5). The 5-HT-induced hyperpolarization was significantly reduced when acutely changing from SLS to malate-SLS (-53 ± 3 mV, n = 5, p = 0.0001, paired two-sided Student’s t-test). On the other hand, when stored continuously in malate-SLS, wash-in of 1 mM 5-HT caused E_m_ to only very slightly hyperpolarize to ∼-50 mV ± 4 mV (n = 5) after 3 min, which renders malate-SLS unsuited for a whole range of experiments. When stored continuously in pyruvate-SLS, 5-HT caused a typical hyperpolarization to ∼-56 mV ± 5 mV (n = 4) on day 1. This 5-HT-induced hyperpolarization in pyruvate-SLS was almost unchanged up to day 3.

In addition to changes in the E_m_ of neurons and glia cells and the AP generation of neurons, isolated leech segmental ganglia also undergo a marked change in the transparency of the ganglion capsule under *ex vivo* conditions. After two or three days in SLS or glucose-SLS, the ganglia appear hazy and blurred (Fig. 3). In pyruvate-SLS this was delayed by about one day and in malate-SLS by 8 days. Apparently, this morphological change corresponds well with the electrophysiological changes.

Despite its moderate effect on extending the viability of leech neurons when compared to malate, pyruvate seemed the best compromise for our tests in acute brain slice preparations in rodents due to absence of obvious changes to tested active (5-HT, AP generation) or passive (E_m_) electrophysiological characteristics.

### **b)** A modified ACSF allows to extend cell viability in acute brain slices to 36 h and beyond

#### Incubation and staining

The UKM chamber allowed stable incubation by deeper wells and circular flow of the superfusion medium (see methods, Fig. 4A-C). The translucent acrylic body with a temperature sensor can be placed into the black heating element for heating to 32 °C or moved to cold storage. Single acute slices from this chamber were moved to a 5 ml staining chamber filled with standard-ACSF and fluorescent dyes from a viability imaging assay: Living cells were stained by Calcein-AM (requiring esterase activity, Fig. 4D, green), dead cells were stained by BOBO-3 iodide (a dsDNA-binding dye, Fig. 4D, red, see methods for details).

**Fig. 4:**
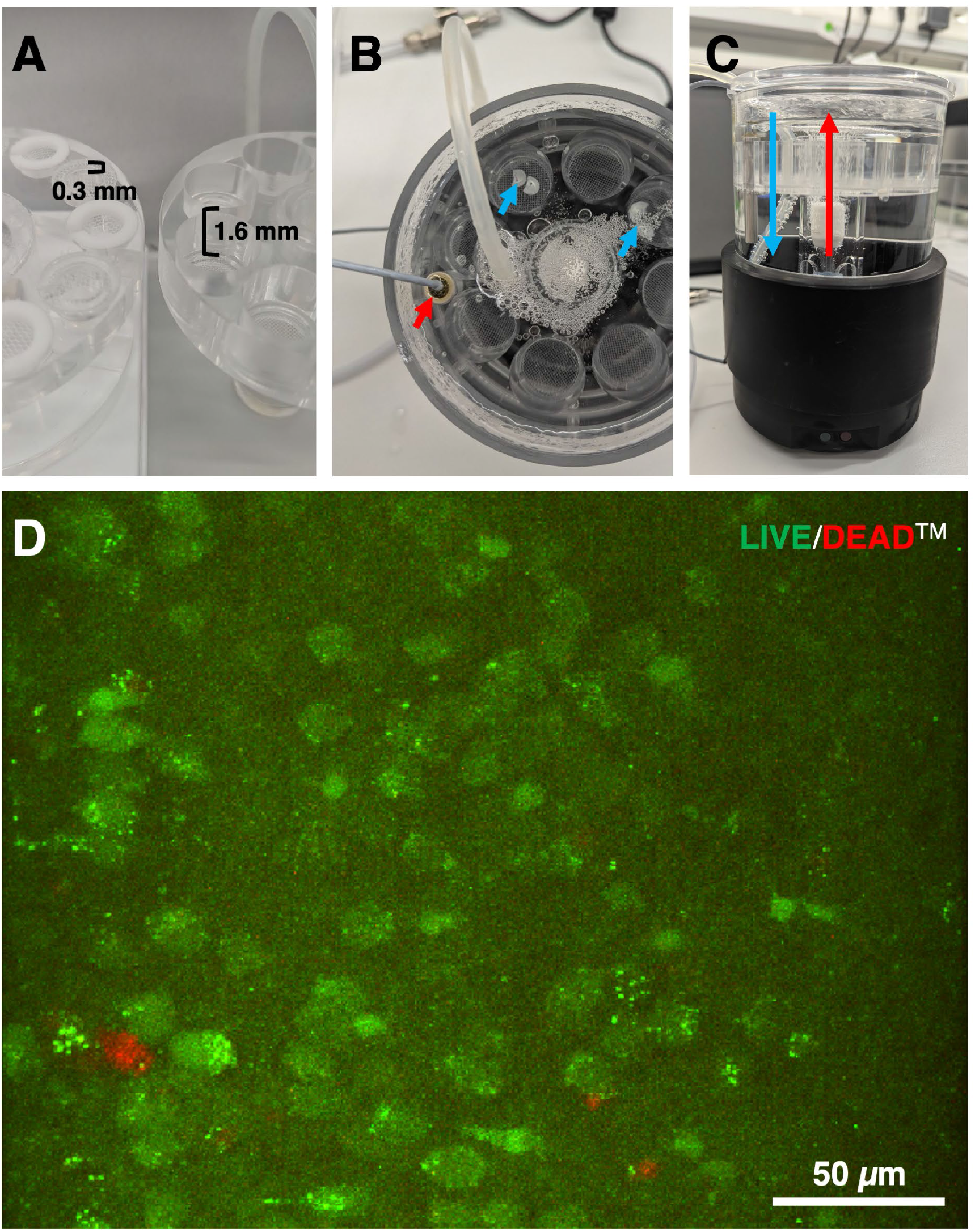
Slice incubation. **A)** Comparison of incubation chamber insets. The inset of the commercially available 12-well chamber (left) had 0.3 cm well depth. Our 8-well chamber (right) was optimized to allow moving it to cold storage by 1.6 cm deep wells. Both were used for gassing with carbogen (95% O2, 5% CO2). **B)** The translucent chamber with a temperature sensor (red arrow) can be placed into a heating element for heating to 32 °C (see also **C**). Slices (blue arrows) were placed on the mesh. **C)** Slices were held firmly in place by circular flow of the superfusion medium from the heating element (black), via a gas bubbler (white), through a vertical channel (red arrow), through the deep wells, over the slices on the mesh and back down (blue arrow) to the heating element. **D)** Cells from mPFC Layer 2/3 on the day 2. Image is a composite of 40 optical slices of 1 μm thickness and thus represents a 40 μm z-stack. Living cells were stained by Calcein-AM (requiring esterase activity, green), dead cells were stained by BOBO-3 iodide (a dsDNA-binding dye requiring a compromised plasma membrane to reach the nucleus, red).

### Imaging essay

Optical sections of 30 or 40 μm thickness (z-stacks) of Layer 2/3 mPFC neurons were analysed (Figs 4-6). These show various degrees of deterioration up to day 4 *ex vivo* (days 1-3 see Fig. 5, day 4 see Fig. 6). In slices that had been kept in pyruvate-ACSF, healthy (green) neurons were markedly more numerous than in slices kept in standard-ACSF for the same duration (Fig. 5). Complete disintegration of the cell membrane, with unhindered staining of the nucleus by BOBO-3 iodide was first observed sparsely on the third day in slices that had been incubated in pyruvate-ACSF. In standard-ACSF, this occurred already on the second day (Fig. 5).

**Fig. 5:**
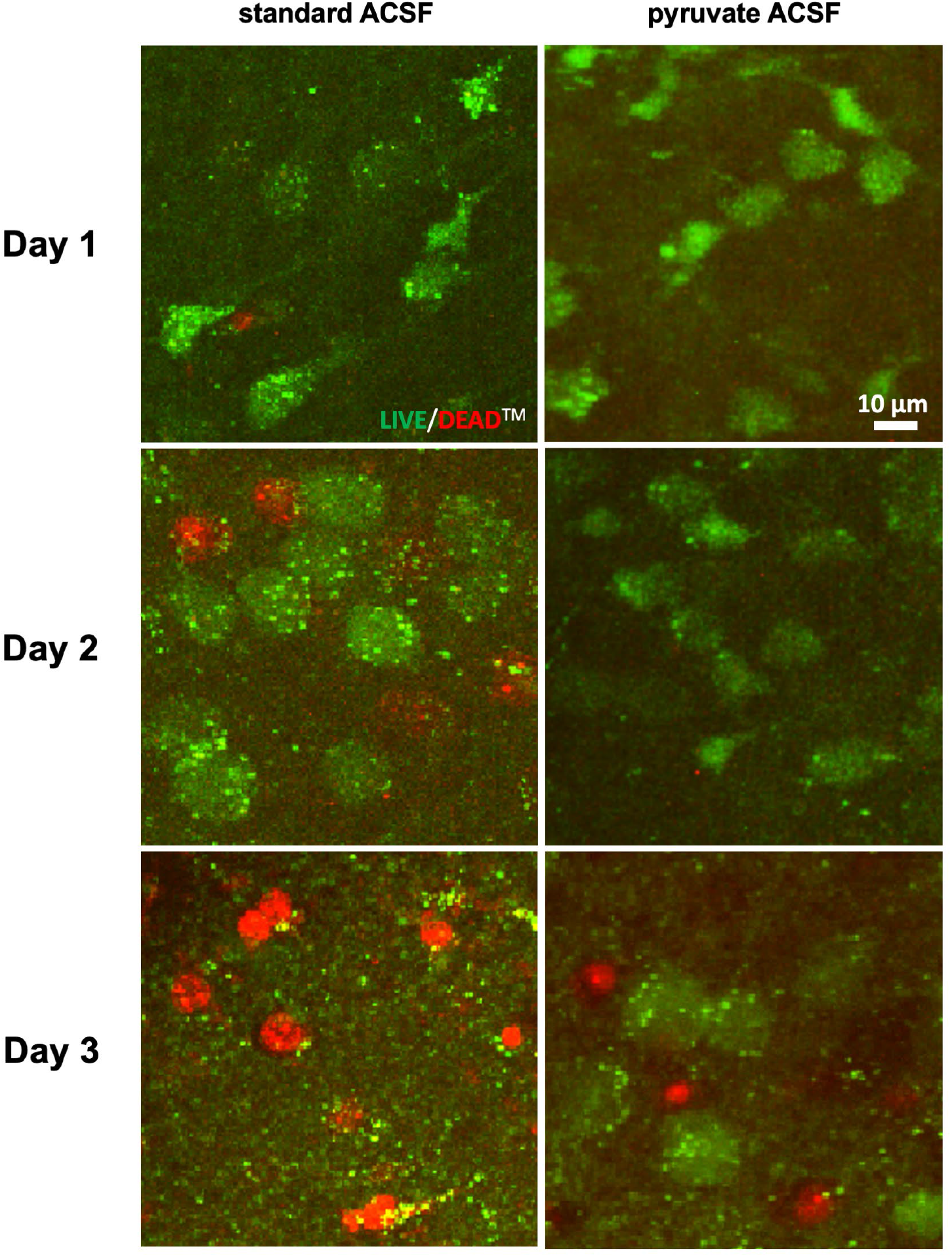
Live/Dead cell imaging essay. Live/Dead Imaging Essay as in Fig. 4D. Comparison of 30 μm z-stacks to assess cell viability in acute slices up to the third day after the slicing procedure. Slices were kept in standard-ACSF (left images) or pyruvate-ACSF (right images).

However, even cells that were stained green must be considered unhealthy when their diameter was markedly increased (Fig. 5, day 2 in standard-ACSF, day 3 in pyruvate-ACSF), which indicates cell swelling (bloating). Similarly, cells must be assumed unhealthy when they no longer retain their morphology (rounding), when their nucleus is well visible (condensation), or vacuole formation has occurred. Such cells could not be recorded with the patch-clamp technique, since they burst or disintegrated when touched by a glass electrode.

The percentage of viable cells in slices that had been incubated in pyruvate-ACSF remained at 90% during the first two days after slicing, followed by a drop to 60% (Fig. 6A). Slices incubated in standard-ACSF showed a noticeable decrease in cell viability to 80% already on the first day, with a further decrease to 60% on the second day.

**Fig. 6:**
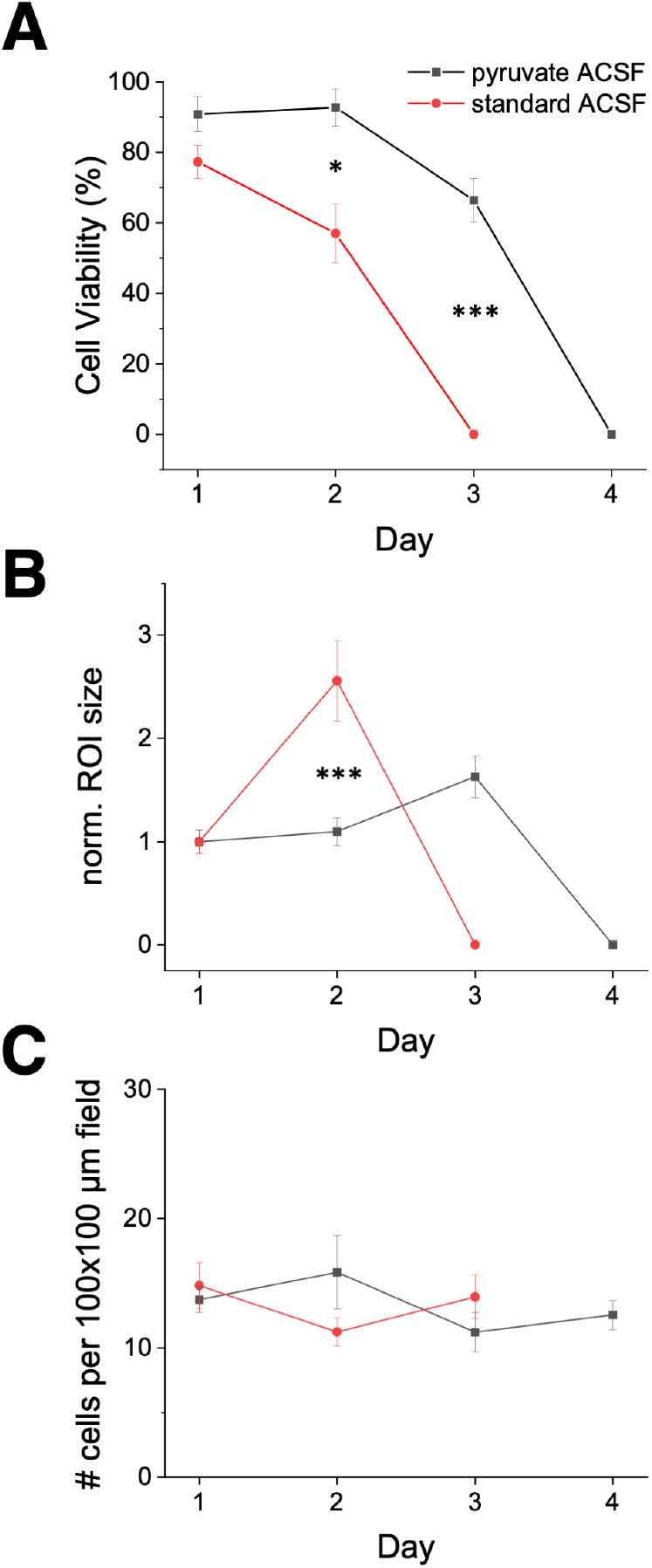
Improved cell viability. Example data from layer 2/3 mPFC cells (see Fig. 5). **A)** Total cell viability in % of all marked cells. **B)** Cell size as normalized ROI size of viable cells depended on time after preparation and on incubation solution. **C)** Cell density was largely unchanged. Between the standard- and pyruvate-ACSF groups, pairwise differences were analyzed using the Mann-Whitney test with Holm-adjusted p-values, *p < 0.05, ***p < 0.001.

The normalized ROI size of cells in field of view in slices that had been kept in pyruvate-ACSF remained unchanged during the first two days and then increased 1.5-fold, while the cell diameters in slices that had been kept in standard-ACSF increased 2.5-fold already by the second day (Fig. 6B).

The total number of cells contained within an area of 0.01 mm^2^ (and 30 μm thickness) remained unchanged at ∼13 cells over the four days post brain slice preparation (Fig. 6C).

Therefore, the use of pyruvate-ACSF allows acute brain slices to be used at least one day longer when compared to a classical incubation protocol (Fig. 5), while the size and viability of such cells on the second day remain on the same level as immediately after the slicing procedure (Fig. 6A, B).

### Electrophysiological characterization

Several electrophysiological properties of neurons in mice of different age groups were evaluated in four brain areas. The areas were chosen to be at different distances from the edge of the coronal brain slice and to have differing neuron and fibre densities. Since the recordings were only used for assessing electrophysiological characteristics for the present study, we generally attempted to continue to use them for further experiments for other, as yet unpublished studies. As this occurred afterwards, such continued use had no impact on this study, but maximized the use of animals in accordance with the 3R principle.

Based on our results from leech neurons, we focussed on comparing the properties of neurons on the first day (i.e. within the first ∼12 h after the slicing procedure) with properties of neurons on the second day (i.e. within ∼24 to 36 h after slicing). For a subset of experiments, we attempted to obtain data from neurons on the third and fourth day (i.e. more than 48 h after slicing). We compared what we consider to be a representative collection of biophysical passive membrane properties (Fig. 7) and active firing properties (Fig. 8, see Methods and Tables 3-7). We briefly describe their presumed value for assessing cell viability in this particular context in the following:

**Fig. 7:**
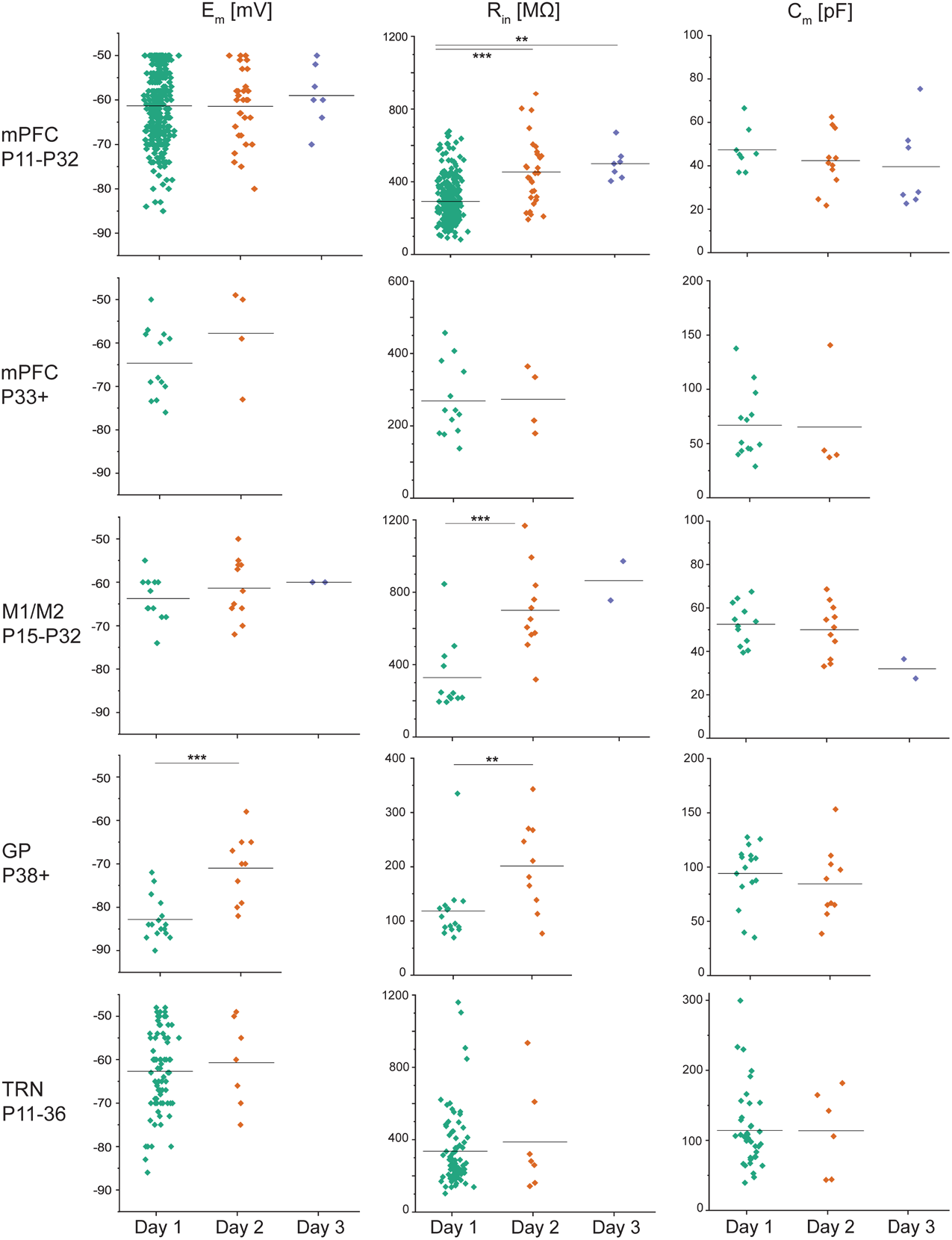
Biophysical, passive electrophysiological characteristics. Biophysical properties of neurons (E_m_, R_in_, C_m_) recorded on the day of the slice preparation (day 1) and the next days (day 2 and sometimes day 3) in L1 of the mPFC, L1 of the M1/M2, rostral TRN, and GP. The age is given in postnatal days. Whole-cell patch-clamp recordings were performed in standard-ACSF at 22 °C. Values from individual neurons are shown as a scatter plot, black lines indicate the mean. Statistical comparisons: Mann– Whitney U test. For two-group comparisons: *p* < 0.05 (*\*), p < 0.01 (**), p < 0.001 (\*\*\**). For three groups comparisons: *p* < 0.0167 (\**), p < 0.0033 (**), p < 0.00033 (\*\*\**). All electrophysiological properties are listed in Tables 3-7.

**Fig. 8:**
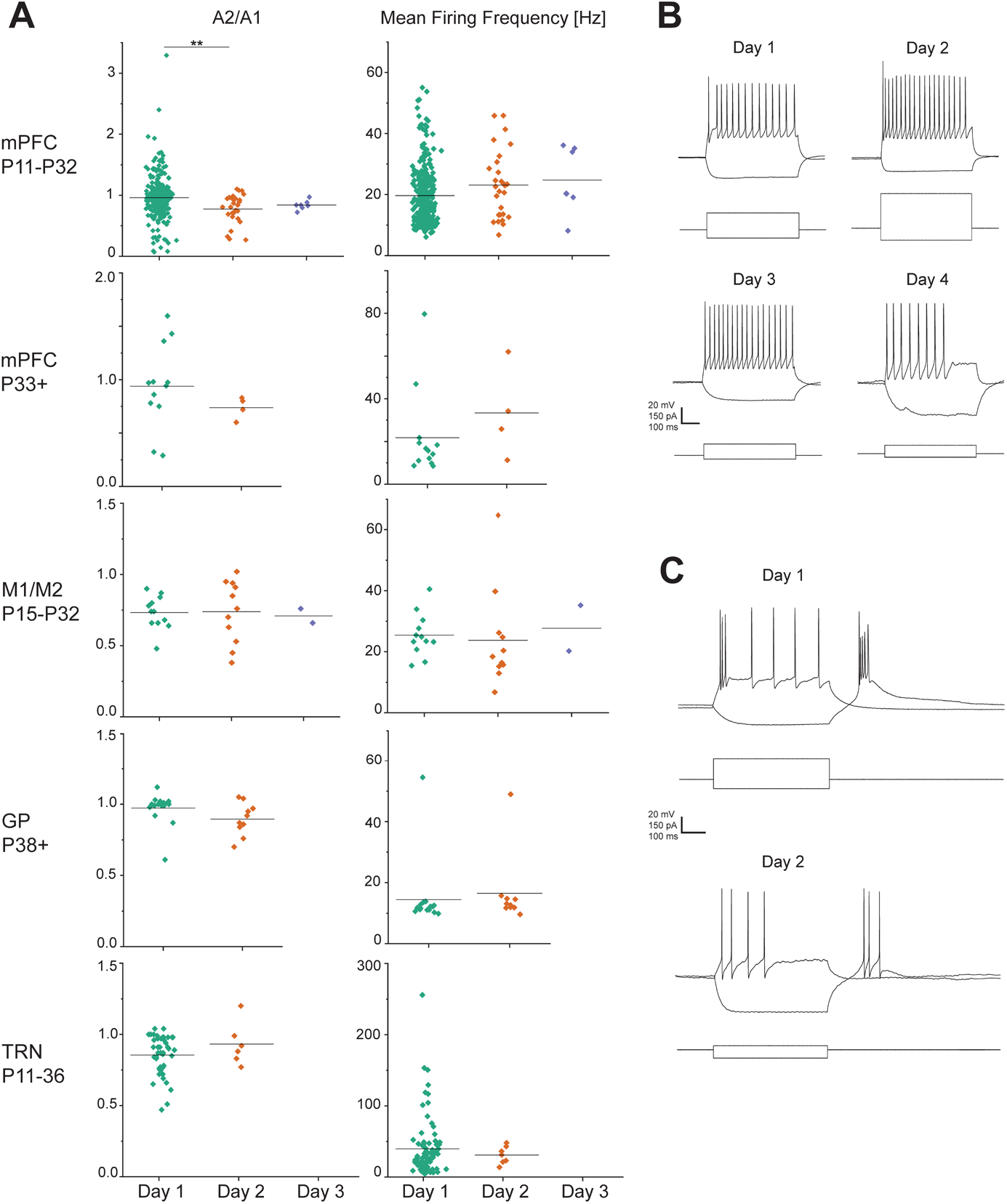
Active electrophysiological characteristics. **A)** Active response properties of the same neurons as in Fig. 7. Action potential amplitude ratio (A2/A1) and the mean frequency of action potential generation (mean firing frequency) at near-threshold current injection (min. four APs, see methods) were calculated from recordings obtained on the day of the slicing procedure (day 1) or the next days (day 2 and sometimes day 3) in L1 of the mPFC, L1 of the M1/M2, TRN, and GP. Age is given in postnatal days. Whole-cell patch-clamp recordings were performed in standard-ACSF at 22 °C. Values from individual neurons are shown as a scatter plot, black lines indicate the mean. Statistical comparisons: Mann– Whitney U test. For two-group comparisons: *p* < 0.05 (*\*), p < 0.01 (**), p < 0.001 (\*\*\**). For three groups comparisons: *p* < 0.0167 (\**), p < 0.0033 (**), p < 0.00033 (\*\*\**). All electrophysiological properties are listed in Tables 3-7. **B)** Current clamp example traces obtained from L1 mPFC neurons from day 1 to day 3 and from L 2/3 on day 4. Hyperpolarizing or depolarizing current steps were chosen to show tonic firing in the depolarizing step and a hyperpolarization to at least -80 mV in the hyperpolarizing step. Traces under day 1 were from an L1 interneuron with the following properties: E_m_ = -70 mV, R_in_ = 444 MΩ, C_m_ = 45 pF; day 2: L1 interneuron, E_m_ = -63 mV, R_in_ = 252 MΩ, C_m_ = 30 pF; day 3: L1 interneuron, E_m_ = -50 mV, R_in_ = 670 MΩ, C_m_ = 25 pF; Day 4: L2/3 neuron, presumably an interneuron, E_m_ = -50 mV, R_in_ = 1036 MΩ, C_m_ = 36 pF. **C)** Current clamp example traces obtained from TRN neurons from the day of the slicing procedure and the day after. Day 1: E_m_ = -66 mV, R_in_ = 319 MΩ, C_m_ = 156 pF; day 2: E_m_ = -61 mV, R_in_ = 830 MΩ, C_m_ = 44 pF. Current injection was chosen to show tonic firing in the depolarizing step and rebound bursting after the hyperpolarizing step.

**Table 3:** Young neurons of the medial prefrontal cortex. Electrophysiological properties of mPFC neurons of young animals (P11 to P32) recorded from cortical layer 1 on the day of slicing (day 1) or the following days (day 2, day 3). See text for details. Age of animals is given as a range in postnatal days. The number of animals is given as N. Values are mean ± SEM of n neurons.

| mPFC (young) | Day 1 (N = 64)<br>Age 11 - 32 | Day 2 (N = 11)<br>Age 11 - 32 | Day 3 (N = 1)<br>Age 22 |
| --- | --- | --- | --- |
| $E_m$ [mV] | $-61 \pm 0.4$ (n = 329) | $-61 \pm 1$ (n = 30) | $-59 \pm 2$ (n = 7) |
| $R_{in}$ [MΩ] | $292 \pm 7$ (n = 305) | $454 \pm 34$ (n = 30) *** | $500 \pm 34$ (n = 7) ** |
| $C_m$ [pF] | $47 \pm 4$ (n = 8) | $44 \pm 6$ (n = 14) | $38 \pm 7$ (n = 7) |
| $C_{in}$ [pF] | $21 \pm 1$ (n = 316) | $22 \pm 2$ (n = 27) | $19 \pm 2$ (n = 7) |
| $\tau$ [ms] | $14 \pm 2$ (n = 8) | $18 \pm 1$ (n = 11) | $19 \pm 3$ (n = 7) |
| Initial firing frequency [Hz] | $25 \pm 1$ (n = 325) | $36 \pm 5$ (n = 30) | $31 \pm 5$ (n = 7) |
| Mean firing frequency [Hz] | 20 ± 0.5 (n = 315) | 23 ± 2 (n = 28) | 25 ± 4 (n = 7) |
| Late firing frequency [Hz] | 17 ± 0.5 (n = 315) | 20 ± 2 (n = 28) | 21 ± 3 (n = 7) |
| AP amplitude ratio (A2/A1) | 0.96 ± 0.02 (n = 315) | 0.77 ± 0.04 (n = 28) ** | 0.8 ± 0.03 (n = 7) |
| AP half-width [ms] | 1.6 ± 0.04 (n = 326) | 1.5 ± 0.1 (n = 30) | 1.5 ± 0.2 (n = 7) |

The **resting membrane potential (E_m_)** allows to assess ion gradients, electromotive forces, ion channel states/membrane permeability, as well as the state of ATP-dependent transport systems, which may be of particular concern during an *ex vivo* incubation. Shifts in E_m_ may indicate changes in the activity of ion channels and transporters (on an expanded time scale).

The **input resistance (R_in_)** determines the impact of an input current on the membrane potential (ΔV/ΔI). Thus, R_in_ determines the amplitude of a neuron’s passive response to incoming signals, such as excitatory or inhibitory postsynaptic currents or any other type of induced transmembrane charge movement. The higher R_in_, the stronger the effect of such charge movements on the cell’s E_m_. Reducing R_in_ as a cellular mechanism can reduce the weight of an input (shunting inhibition). Changes in R_in_ reflect changes in the number of open or active ion channels.

The **membrane capacitance (C_m_)** is proportional to the surface area of the cell membrane. It reflects the ability of the cell to temporarily store charges and thereby, together with R_in_, determines how rapidly E_m_ may change. Thus, the product of R_in_ and C_m_ is **membrane time constant (**τ). It determines the time passed until the membrane reaches ∼63% (1-1/e) of a voltage change at any given current injection.

The frequency of action potential generation, often referred to as **spike frequency**, is dependent on many factors, involving E_m_, ion channel states, [Ca^2+^]_i_ which in turn defines the state of Ca^2+^-dependent Cl^-^and K^+^ channels [62,63]. This was assessed in three ways: the **initial spike frequency** is the unaltered and immediate frequency of just the first two action potentials. While it may be subject to fast adaptation mechanisms, such as an altered availability of Na_v_ channels between the first and the second action potential, Ca^2+^-dependent spike frequency adaptation should not yet play a role [63,64]. The **late spike frequency**, however, can elucidate spike frequency adaptation [65]. And finally, the **mean spike frequency** may allow a more general comparison of all factors involved and may represent the overall responsiveness of a neuron.

The **First AP half-width** or full-width-at-half-maximum (**FWHM**) is dependent on the reaction speed of the ion channels involved in the Hodgkin cycle, but also on C_m_ and R_in_.

The long-term maintenance of the ionic gradients of Na^+^ and K^+^ across the neuronal membrane requires the activity of the Na^+^/K^+^ ATPase. A lack of energy sources will thus cause the maintenance to fail and will have effects on, among others, E_m_ and action potential amplitude. This can be assessed by the **action potential (AP) amplitude ratio**, where the amplitude of an action potential is divided by the amplitude of its antecedent AP.

TRN neurons are characterized by their ability to generate low-threshold bursts, also termed low-threshold spikes (LTS). These are mediated by T-type Ca^2+^ channels (Ca_v_ 3.x) that produce a slow (∼200 ms) depolarizing envelope crowned by fast Na^+^/K^+^ action potentials at very high frequencies [59,61,66]. The **frequency** with which such bursts occur, the **width** of the Ca^+^ envelope, and the number and frequency of **spikes in a burst** were analyzed [cf. 59].

Thus, the electrophysiological characteristics of neurons in mPFC (layer 1, separated in young animals and animals older than P33), M1/M2 (layer 1), GP, and the rostral TRN, obtained at day 1, day 2, and sometimes day 3 after the slicing procedure, were analyzed (Tables 3 to 7).

### mPFC

The cortical layer 1 (L1) consists mainly of distal apical dendrites of pyramidal neurons from the underlying cortical layers, thalamocortical inputs, and a sparse population of two types of GABAergic interneurons [67–69], the single bouquet cells (SBC) and (elongated) neurogliaform cells (NGFC, ENGC) [70–72]. We focused on the properties of cells in the L1 of infralimbic and prelimbic mPFC, positioned at the midline, and the motor cortex (M1 and M2), positioned at the outer edge of the brain slice. Electrophysiological properties were assessed for juvenile animals aged P11-32 (Table 3, Figs 7-8) and young adult animals (P33+, Table 4, Figs 7-8).

**Table 4:**
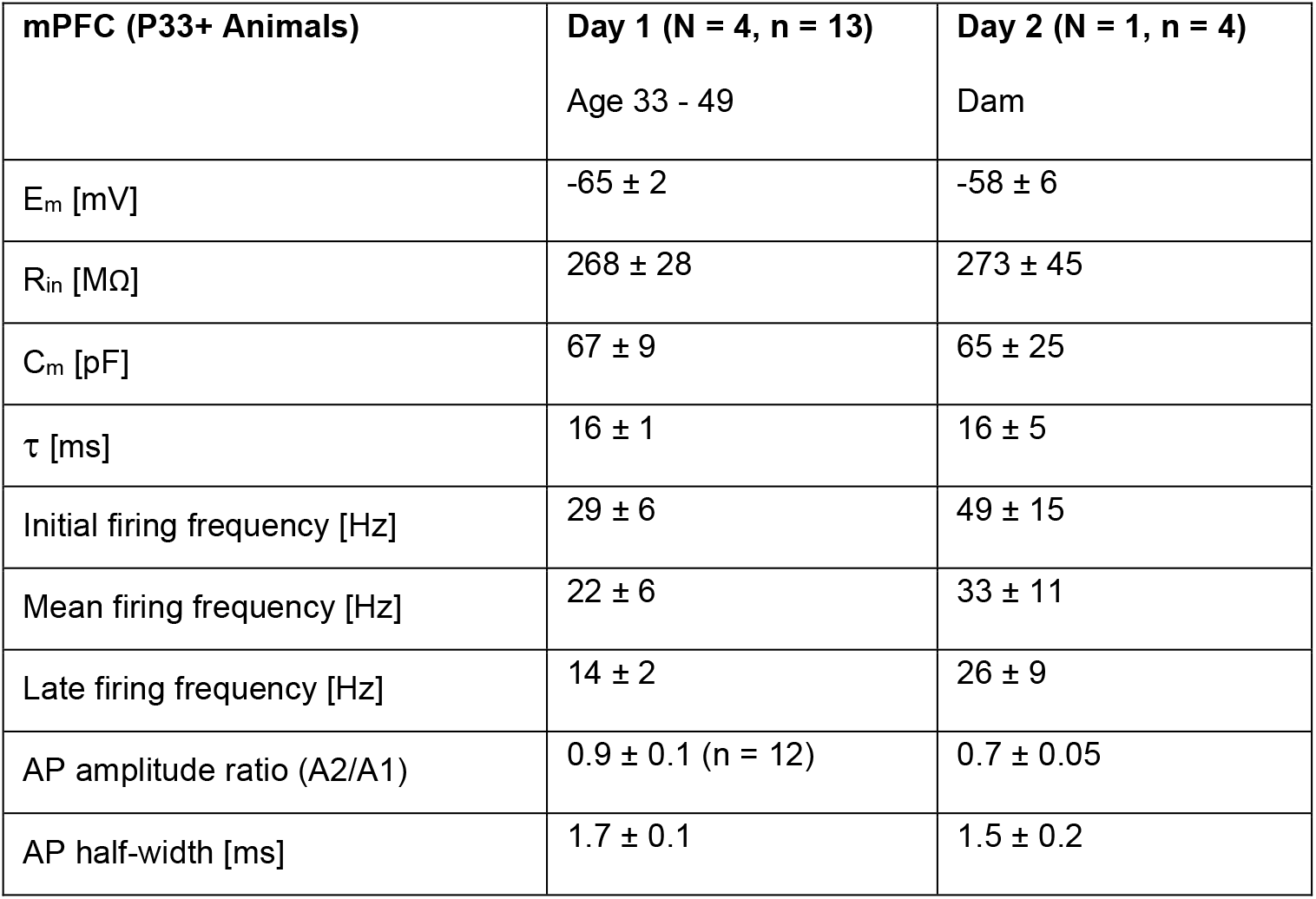
Older neurons of the medial prefrontal cortex. Electrophysiological properties of mPFC neurons of animals aged P33 and older, recorded from cortical layer 1 on the day of slicing (day 1) or the following day (day 2). See text for details. Age of animals is given as a range in postnatal days. The number of animals is given as N. Values are mean ± SEM of n neurons.

Neurons from juvenile animals retained all electrophysiological characteristics on day 2 and day 3, except R_in_ and the AP amplitude ratio. Key values were consistent with previously published data [73–75]. The AP amplitude ratio changed from 0.96 ± 0.02 on day 1 to 0.77 ± 0.04 on day 2 and remained constant at 0.8 ± 0.03 on day 3. Similarly, on day 2, R_in_ increases quite strongly from 292 ± 7 MΩ to 454 ± 34 MΩ and remains on a similar level on day 3. For older animals, neurons in the mPFC had slightly altered parameters, albeit none were significantly different (Table 4). The observed increases in firing frequency (upon current injection, see methods) and R_in_ are unlikely to be a sign of degradation (see discussion).

### M1/M2

The neurons tested in layer 1 of the motor cortex (M1/M2) showed a very strong and significant increase in R_in_ from 328 ± 56 MΩ to 700 ± 71 MΩ, a significant increase of τ by a factor of more than two (from 17 ± 3 ms to 36 ± 4 ms), and a slight increase of the FWHM from 1.4 ± 0.1 ms to 1.8 ± 0.1 ms (see Table 5, Figs 7-8). Since C_m_ remains unchanged, these changes are unlikely to indicate a change in membrane surface area but may indicate a change in the voltage dependency of activation of ion channels. However, since FWHM is unaltered, this should exclude ion channels that determine the shape of the action potential (see discussion).

**Table 5:** Neurons of the motor cortex. Electrophysiological properties of M1/M2 neurons recorded from cortical layer 1 on the day of slicing (day 1) or the following days (day 2, day 3). See text for details. Age of animals is given as postnatal days. The number of animals is given as N. Values are mean ± SEM of n neurons.

| **M1/M2** | **Day 1**<br>**(N = 2, n = 12)**<br>Age 28, 32 | **Day 2**<br>**(N = 2, n = 11)**<br>Age 15, 17 | **Day 3**<br>**(N = 1, n = 2)**<br>Age 17 |
| --- | --- | --- | --- |
| $E_m$ [mV] | $-64 \pm 1$ | $-61 \pm 2$ | -60 |
| $R_{in}$ [$\text{M}\Omega$] | $328 \pm 56$ | $700 \pm 71$ *** | 864 |
| $C_m$ [pF] | $52 \pm 3$ | $50 \pm 4$ | 32 |
| $C_{in}$ [pF] | $19 \pm 2$ | $14 \pm 2$ | 21 |
| $\tau$ [ms] | $17 \pm 3$ | $36 \pm 4$ ** | 28 |
| Initial firing frequency [Hz] | $31 \pm 7$ | $31 \pm 8$ | 36 |
| Mean firing frequency [Hz] | $25 \pm 2$ | $24 \pm 5$ | 28 |
| Late firing frequency [Hz] | $21 \pm 1$ | $23 \pm 5$ | 23 |
| AP amplitude ratio (A2/A1) | $0.73 \pm 0.03$ | $0.74 \pm 0.07$ | 0.71 |
| AP half-width [ms] | $1.4 \pm 0.1$ | $1.8 \pm 0.1$ ** | 1.7 |

### GP

The globus pallidus (GP) is a nucleus of the basal ganglia which is believed to play a central role in Parkinson’s Disease [76]. It consists mainly of GABAergic neurons of two subtypes (prototypic and arkypallidal cells, [77]) that preferentially project to the subthalamic nucleus and the striatum [77]. We tested our preparation on these neurons and assessed their electrophysiological properties in adult animals (P33+) within 36 h after the slicing procedure (Table 6, Figs 7-8). Although R_in_ increased and E_m_ depolarized on day 2 compared to day 1, values remained well within previously reported ranges [78–81]. The limited variability of E_m_ and R_in_ observed on day 1 in recordings from a single animal (Figs 7-8) suggests that the neurons may represent a single neuronal subtype or reflect subtle differences in experimental conditions (see discussion).

**Table 6:** Neurons of the globus pallidus. Electrophysiological properties of GP neurons on the day of slicing (day 1) or the following day (day 2). See text for details. Age of animals is given as postnatal days. The number of animals is given as N. Values are mean ± SEM of n neurons.

| GP | Day 1 (N = 1, n = 16) | Day 2 (N = 2, n = 10) |
| --- | --- | --- |
|  | Age 38 | Age 38, dam |
| $E_m$ [mV] | $-83 \pm 1$ | $-71 \pm 2$ *** |
| $R_{in}$ [MΩ] | $118 \pm 15$ | $208 \pm 31$ ** |
| $C_m$ [pF] | $94 \pm 7$ | $84 \pm 10$ |
| $\tau$ [ms] | $10 \pm 0.8$ | $17 \pm 3$ |
| Initial firing frequency [Hz] | $19 \pm 4$ | $16 \pm 2$ |
| Mean firing frequency [Hz] | $14 \pm 3$ | $16 \pm 4$ |
| Late firing frequency [Hz] | $13 \pm 2$ | $14 \pm 3$ |
| AP amplitude ratio (A2/A1) | $0.97 \pm 0.03$ | $0.90 \pm 0.05$ |
| AP half-width [ms] | $1.8 \pm 0.1$ | $1.9 \pm 0.2$ |

### TRN

The thalamic reticular nucleus (TRN) is a thin layer of exclusively GABAergic neurons surrounding the dorsal thalamus [61,82–84]. Because the TRN is characterized by a high density of fibres, both passing and terminating, from both thalamocortical and corticothalamic projections, recording from TRN neurons always poses a technical challenge and therefore, our previously published studies used slices from younger animals [59,85–87], when projections are less dense but already exhibit adult features [88,89]. In the present study, despite considerable improvements, even on day 1, and increasingly so on subsequent days, obtaining a stable recording remained a challenge and only seven neurons were recorded from day 2. Their electrophysiological properties did not differ from TRN neurons recorded on day 1 (Table 7, Figs 7-8) and were similar to previously published values [59,90–93]. However, while our preparation procedure may improve recordings (or the likelihood of obtaining recordings) from the TRN on day 1, TRN neurons should not be expected to be easily accessible on day 2.

**Table 7:** Neurons of the thalamic reticular nucleus. Electrophysiological properties of TRN neurons recorded on the day of slicing (day 1) or the next day (day 2). See text for details. Age of animals is given as a range in postnatal days. The number of animals is given as N. Values are mean ± SEM of n neurons.

| TRN | Day 1 (N = 40)<br>Age 11-36 | Day 2 (N = 5)<br>Age 12 - 28 |
| --- | --- | --- |
| $E_m$ [mV] | $-63 \pm 1$ (n = 81) | $-61 \pm 4$ (n = 7) |
| $R_{in}$ [MΩ] | $336 \pm 22$ (n = 80) | $387 \pm 108$ (n = 7) |
| $C_m$ [pF] | $115 \pm 8$ (n = 45) | $114 \pm 24$ (n = 6) |
| $C_{in}$ [pF] | $51 \pm 4$ (n = 81) | $34 \pm 5$ (n = 7) |
| $\tau$ [ms] | $41 \pm 3$ (n = 46) | $37 \pm 6$ (n = 6) |
| Initial firing frequency [Hz] | $69 \pm 10$ (n = 46) | $45 \pm 26$ (n = 6) |
| Mean firing frequency [Hz] | $40 \pm 4$ (n = 81) | $31 \pm 5$ (n = 7) |
| Late firing frequency [Hz] | $25 \pm 5$ (n = 46) | $23 \pm 12$ (n = 6) |
| AP amplitude ratio (A2/A1) | 0.86 ± 0.02 (n = 46) | 0.93 ± 0.06 (n = 6) |
| AP half-width [ms] | 1.20 ± 0.07 (n = 46) | 1.14 ± 0.16 (n = 6) |
| LTS propensity [0-1] | 0.7 (n = 81) | 0.6 (n = 7) |
| Oscillations [0-1] | 0.5 (n = 59) | 0.5 (n = 4) |
| Width of LTS [ms] | 236 ± 8 (n = 59) | 236 ± 10 (n = 4) |
| Number of spikes per LTS | 4.1 ± 0.3 (n = 59) | 3 ± 1 (n = 4) |
| Initial Firing Frequency in RB [Hz] | 153 ± 10 (n = 28) | 101 ± 29 (n = 3) |
| Firing Frequency in RB [Hz] | 116 ± 6 (n = 52) | 81 ± 28 (n = 3) |
| Amplitude of the 1<sup>st</sup> spike in RB [mV] | 65 ± 2 (n = 59) | 55 ± 9 (n = 4) |
| AP amplitude ratio (A2/A1) in RB | 0.6 ± 0.04 (n = 28) | 0.92 ± 0.02 (n = 3) |

## 4. Discussion

### Aim

The European Union has specific legislation on the use of animals in science. Since EU-Directive 2010/63/EU came into effect, the 3R principle is explicitly established in EU member states, requiring experimenters to *reduce*, *refine*, or *replace* animal use in research. While the 3R principle was coined in 1959 by Russel & Burch [94], the concept is much older and was, for instance, the subject of a letter by Marshall Hall to the editor of the journal Lancet in 1847 [95]. Today, the ethical framework for improving animal welfare continues to evolve and is widely adopted [1].

Acute brain slice preparations allow to experimentally assess neuronal function without ongoing animal experimentation and, therefore, fulfill the criterion of *refining* by reducing animal suffering. Our aim was to extend brain slice viability to additionally allow to *reduce* the number of animals used, which we achieved with the described preparation, slicing, and incubation procedure. We were able to store brain slices overnight for next day use, thus considerably reducing the number of animals we required.

### Approach

It was previously reported that modified preparatory solutions based on NMDG and incubation solutions containing e.g. L-ascorbic acid, thiourea, N-acetylcysteine, and pyruvate (see methods) improved the "general appearance of neurons" and improved the "ease of patch clamp recording" [9], an observation that we can verify. Interestingly, it was also reported that slices could be "maintained in a healthy state for extended time periods" [9] without defining what exactly could be expected. Our superfusion media, and our method of slice incubation, are all based on the idea that they should provide the following key functions:

1. Providing sufficient oxygenation of the neuronal tissue and thus preventing hypoxia. This is generally achieved by gassing with carbogen, which also ensures a stable pH in combination with sodium bicarbonate (NaHCO_3_). But the influence of metabolic rate and temperature on oxygen requirements should not be neglected: experimental procedures often involve processes that increase metabolic rate (neurotransmitter release, second messenger signalling, Ca^2+^-entry and -extrusion, excitation) and "the capacity of oxygen delivery matches full aerobic scope only within the thermal optimum" [96]. Thus, high oxygen supply or high superfusion rates are beneficial for neuron survival, but a low metabolic activity (e.g. by reduction of temperature during our cold storage and by removal of Na^+^ in our recovery-ACSF) is also a viable strategy and reflects the use of hypothermia as a neuroprotective measure in patients with brain hypoxia [97]. The short-term use of NMDG to replace Na^+^ was consistently shown to have neuroprotective effects [15,16,50].
2. Removal or decomposition of metabolic end products. This either requires constant exchange of the holding medium, filtering (as reported previously [21–24]), or a large enough volume to limit the accumulation of detritus, as in our holding chamber (Fig. 4).
3. Supply of molecules for the cellular energy metabolism. So far, the primary compound considered for this was Glucose (D-Glucose or Dextrose). In a series of small thesis projects using leech ganglia (supervised by P. H.), the impact of anaplerotic components (anaplerosis by ingestion of organic acids, [98]) was tested and we present some of the results here for the first time (except [32]). Surprisingly, these experiments uniformly revealed that glucose had no impact whatsoever on the viability of neurons and neuropile glia cells in leech segmental ganglia (Fig. 2-3). Instead, pyruvate and other components of the tricarboxylic acid cycle (TCA cycle), were found to be essential for maintaining viability, with Malate being the most effective. Pyruvate was repeatedly found to improve the preparation of brain slices [16,50] and it plays a major role in the "Astrocyte-to-Neuron Lactate Shuttle" (ANLS) hypothesis [99]. This hypothesis suggests that, in the human brain, lactate produced by astrocytes is transferred to neurons, where it is converted to pyruvate for aerobic energy production via the TCA cycle [100]. In support of this, 2.5 h after slicing, lactate levels started to gradually decrease in brain tissue, while the pyruvate levels started to increase over the same time period [101], likely affecting neuronal function. Supplementing pyruvate via the holding solution would counteract this and keep lactate levels stable, while providing an all but unlimited supply of pyruvate to the TCA cycle of neurons and glia cells.
4. Minimizing mechanical damage. The use of an incubation chamber with deep wells helps prevent the slices from swirling around during transfer of the chamber to cold storage (see methods). In addition, bubbling from the centre of the chamber causes a circular flow of the medium from the centre toward the surface of the medium, and back downward through the wells. This pins the slices to the surface of the supporting mesh (Fig. 4B) and provides a constant stream of freshly oxygenated and buffered medium to the slices [cf. 10]. Additionally, this configuration reduces micromovements induced by the bubbling, or rather by the bursting of bubbles.
5. Reducing bacterial growth. Previously, this was achieved by UV-C treatment [22,23].

Here, we filtered newly made solutions through sterile filters. Additionally, our incubation solution contained ascorbic acid and NAC, which are known to have antibacterial properties [102,103]. Additionally, cold storage will reduce bacterial growth.

Many methods and variations of methods are used for preparing brain slices and the aim of the preparative method is, of course, to produce "healthy" slices. What exactly constitutes healthiness and how to assess it, has been a matter of debate [18]. Four main categories were identified: morphology, electrophysiological characteristics, metabolic characteristics (including phosphorylation of macromolecules), and protein synthesis (including stress markers) [17,18].

Here, we chose to assess I) membrane integrity (Figs 4-5), II) cell morphology (Fig. 6), III) biophysical membrane properties (Fig. 7), active electrophysiological response characteristics (Fig. 8), including possible indicators of intact metabolic functioning (AP half width, AP amplitude ratio) and of the state of modulation by adaptation (late firing frequency, Tables 3-7). We thereby demonstrate that our method allows to extend the viability of acute brain slices to at least 36 h after the slicing procedure with only minor limitations. Although we expanded the range to test even beyond 36 hours after preparation, we suggest to be cautious here. For further optimizing animal use, expanding beyond 36 h is of limited value. Stable recordings from neurons 36 h after preparation is already a major step forward. Further improvements could be valuable for experimental procedures that require longer incubation times.

### Assessing brain slice viability and neuronal health

#### Membrane integrity

Membrane integrity is vital for electrophysiological recordings. Damage to the membrane will cause a breakdown of the neuron’s ion gradients and, thus, of electromotive forces. When this occurs during a whole cell patch-clamp experiment, this will become apparent as a sudden drop in R_in_ and a sudden depolarization of E_m_. Before patching, neurons that have a visibly compromised membrane (bloated cell body and visible nucleus, see day 2 for standard-ACSF and day 3 for pyruvate-ACSF in Fig. 5) will typically be avoided by the experimenter. Therefore, the success rate of obtaining a giga seal or good whole cell access to a neuron is not a reliable parameter for assessing the healthiness of a brain slice preparation, unless the experimenter is forced to choose blindly which cell to target *[-which is just too frustrating to be feasible*].

### Electrophysiological parameters

When determining the health of neurons in patch-clamp experiments, experimenters mostly aim for cells with high R_in_, membrane capacitance, and frequency of AP generation, since these parameters usually indicate an intact and active neuron, efficient signal transmission, and intact brain tissue. For instance, a high firing frequency may indicate an intact energy metabolism [104].

The R_in_ of a neuron at rest is the inverse measure of its leakiness and is governed to a large part by two-pore domain potassium channels [105,106] or by chloride channels such as the CLC-2 [107]. R_in_ crucially affects signal transmission [106,108], and may indicate changes in dendritic access resistance due to compartmentalization [109]. In our experiments on day 2 and thereafter, we observed either an unchanged (older mPFC neurons and TRN neurons, Tables 4 and 7) or an increased R_in_ (young mPFC neurons, M1/M2 neurons, GP neurons, Tables 3, 5 and 6). An increase in R_in_ is not a typical hallmark of decay. On the contrary, dying neurons usually present with a reduced R_in_. This does, in turn, not imply that higher values for R_in_ are evidence of a particularly healthy neuron. There are several possible reasons for an increased R_in_: 1) A neuron’s dendrites may be damaged during the slicing procedure and cause leak currents that lead to a decreased R_in_ on day 1, which could heal during the incubation period. On day 2, the observed increase in R_in_ may then be attributed to this healing process, in which dendrites damaged during the slicing procedure are repaired, causing a reduced current leak; 2) Changes in R_in_ also reflect changes in the number of open or active ion channels, and thus could be an indicator of metabolic breakdown (e.g. lack of phosphorylation of ion channels). 3) Changes in R_in_ may indicate a change in the voltage dependency of activation of ion channels that are active at the voltage ranges we used to determine R_in_. Candidates include K^+^ channels of the M-type (KCNQ/KV7, [110,111]) or hyperpolarization-activated, cyclic nucleotide gated nonselective cation channels (HCN, [105,112]). The latter could be affected via its dependency on cAMP, which may be lacking under the experimental conditions of extended incubation. 4) Increases of R_in_ may be a sign of reduced synaptic input to a neuron [113], which may here have occurred due to gradually degrading synaptic inputs from other neurons in the vicinity of the successfully patched (and obviously healthy) neuron.

However, all of these reasons seem unlikely in light of the largely unchanged firing properties and E_m_ of the tested neurons, as was the case for young mPFC neurons, M1/M2 neurons, and GP neurons, although in young mPFC neurons, A2/A1 slightly decreased (Table 3), in M1/M2 neurons the AP half width slightly increased (Table 5), and in GP neurons E_m_ significantly depolarized (Table 6). The increased AP half width in M1/M2 neurons, and hence the broadening of the AP, is possibly due to the slight, subthreshold depolarization of E_m_ (see Table 5, [114]) or due to the increase in τ but could also suggest altered ion channel contributions (e.g. BK_Ca_^2+^ channel activation, [63,115]), slower kinetics of ion channel opening and closing (such as of K_v_3.4) by changes in phosphorylation [116–118].

Since R_in_ also determines the amplitude of a neuron’s passive response to incoming signals, such as excitatory or inhibitory postsynaptic currents, massive increases in R_in_, as observed for M1/M2 neurons (Table 5) and GP neurons (Table 6) are undesirable in electrophysiological experiments (see above). It remains unclear what caused this.

The resting membrane potential (E_m_) allows to assess ion gradients, electromotive forces, ion channel states and thus membrane permeability, as well as the state of ATP-dependent transport systems (Na^+^/K^+^ ATPase). Thus, shifts in E_m_ may indicate changes in the activity of ion channels and transporters, or a change of ion gradients. Slight depolarizations were registered for older mPFC neurons (albeit not significant) and for GP neurons. In GP neurons, this was accompanied by a significant increase of R_in_ on day 2. The comparably low R_in_ and the markedly hyperpolarized E_m_ of GP neurons on day 1 may indicate the activity of a hyperpolarizing membrane conductance which could be absent on day 2, such as ATP-inhibited K^+^ channels (indicating lower ATP levels on day 1 than on day 2, [119,120]), Ca^2+^ dependent K^+^ channels (indicating higher [Ca^2+^]_i_ on day 1 than on day 2, [59,63,121]), or swelling activated Cl^-^ channels (indicating a cell swelling on day 1 but not on day 2, [3,122]). While it remains unclear whether these changes indicate healing or deterioration, the intact firing properties strongly argue against a substantial damage to GP neurons on day 2.

The frequency of action potential generation was not significantly affected in any of the tested brain areas, similarly to the experiments on leech neurons. Low-threshold bursts with similar properties were observed in TRN neurons of day 1 and day 2.

### L1 interneurons of the mPFC and M1/M2

The cortical layer 1 is unique for its absence of excitatory neurons. Among the dendrites of pyramidal cells from deeper layers is a sparse population of inhibitory interneurons [123]. We here focused on the properties of neurons in the L1 of infralimbic and prelimbic mPFC, positioned at the midline, and the motor cortex (M1 and M2, see below), positioned at the outer edge of the brain slice. The latter neurons are subject to considerable mechanical stress during preparation, due to their exposed position, which has been reported to mimic penetrating brain injury [124]. For the cortical areas investigated here, stable recordings could be obtained up to day 3, i.e. more than 48 h after the slicing procedure, and after twice transferring the slices to overnight cold storage and back.

The significant changes in AP amplitude ratio in young mPFC neurons (Table 3), could indicate a fast and local breakdown of ion gradients, although the stable E_m_ does not support this. Together with the increase in R_in_, this could reflect compartmentalization of the cell, resulting in a smaller contributing cell volume and surface area, and thus a faster exhaustion of available ions, a fast and local breakdown of ion gradients, or gradually ceasing synaptic inputs. The constant C_m_, C_in_, and τ all strongly argue against a reduction of effective surface area. Moreover, the change observed here, despite its statistical significance, is probably too small to have any functional impact. The significant changes in R_in_ are much more physiologically relevant but nevertheless remain within the range observed on day 1. Data in layer 1 of the mPFC for adult animals (P33+, Table 4) is rather sparse. Obtaining data from interneurons in brain slices of older animals is always more of a challenge than in brain slices from younger animals, let alone more than 24 h after the slicing procedure. Thus, the fact that we were able to obtain these recordings must be considered a success. Moreover, it seems that the electrophysiological characteristics of the recorded cells are well within the range of values observed on day 1.

The neurons tested in layer 1 of the motor cortex (M1/M2) showed a very strong and significant increase in R_in_ from 328 ± 56 MΩ to 700 ± 71 MΩ, a significant increase of τ by a factor of more than two (from 17 ± 3 ms to 36 ± 4 ms), and a slight increase of the AP half width from 1.4 ± 0.1 ms to 1.8 ± 0.1 ms. Such an increase in AP half width can cause increased neurotransmitter release at synaptic terminals and thereby modulate synaptic strength [114,125]. The membrane capacitance C_m_ has been used previously as an indicator of cell surface area [3]. Neither C_m_ (determined via τ) nor C_in_ (determined via capacitance neutralization, see methods) indicated an increase of the cell surface area, arguing against the idea of healing.

### TRN

In isolated brain slices, oxygen and nutrient supply is entirely dependent on diffusion from the perfusion medium. In the absence of vascular perfusion, diffusion limitations become more pronounced with increasing cell and fiber density, which may affect metabolic homeostasis and cell viability. Thus, areas with particularly high cell and projection density (such as TRN) could be more prone to hypoxia under standard slice incubation conditions. Nevertheless, for cells that were recorded on day 2 after slicing, there were no differences in intrinsic cell properties, firing and rebound burst activity.

### Lessons from the leech

The central nervous system of the medicinal leech is surrounded and supplied by a hemocoelomic system [39,126]. The blood of leeches contains various inorganic cations and carboxylate anions, with malate being the most prominent component [27,42]. The significance of these carboxylate anions for the function of the leech central nervous system has been studied previously in several theses [e.g. 32]. Malate-supplemented Ringer’s solution was tested on leech ganglia and shown to prolong neuron lifetime from one day to approx. a week.

When transferring isolated leech neurons into cell culture, Leibovitz-15 medium was used, which contains 5 mM pyruvate [33,37,38]. Similar to neurons in our brain slice preparation, the input resistance of leech neurons in culture was also much larger than *in vivo* [127]. We therefore tested a simplified BEL with pyruvate as the only organic anion, revealing that pyruvate similarly extended viability, albeit less so than malate. Moreover, pyruvate had no side effects on the membrane potential (Fig. 2), on active responses (generation of spontaneous action potentials and hyperpolarizing response to 5-HT), or on the transparency of the ganglion capsule (Fig. 3).

The role of glucose as primary fuel source for the brain has been a matter of controversy. It was previously demonstrated that the mammalian brain uses pyruvate as a fuel source [128], and pyruvate can support neuronal viability in the absence of glucose. Mitochondrial pyruvate uptake was identified as a critical node for the metabolic control of neurotransmission in hippocampal terminals. Here, we show that viability of leech ganglia was independent of glucose, but not of pyruvate, supporting this notion. The use of pyruvate was further encouraged by previous publications on a protocol allowing to access adult neurons easily with the patch-clamp technique [16,50]. The effects of 5 mM pyruvate argue in favour of the idea that this substance is metabolized directly in the Krebs cycle by neurons. Moreover, pyruvate is also contained in Leibovitz-15 medium, which allowed to transfer isolated leech neurons into cell culture [33,37,38], even without glia [127] or glucose. For cultivating leech neurons, typically 10 mM glucose were added to Ringer’s solution [7] or Leibovitz-15 medium was used, which contains 30 mM glucose [37]. To test for glucose efficacy to extend the viability of leech neurons, we added 10 mM glucose to SLS. This solution did not extend the viability of leech neurons beyond that in SLS. This raises the long overdue question of why exactly it was added to numerous solutions in the first place.

### Experimental and practical usefulness

We were able to successfully apply the combination of whole-cell patch-clamp recordings and extracellular electrical stimulation using bifurcated stimulus electrodes in TRN to brain slices from day 2 [129]. Moreover, we were able to investigate electrical synaptic plasticity in L1 interneurons of the mPFC using paired whole-cell patch-clamp recordings in brain slices up to 72 h after the slicing procedure [130]. We consider both experiments to be very demanding and thus in need of high-quality brain slices. We suggest that this method could be used to reduce the number of animals in studies involving whole cell patch clamp recording techniques. This may be the case when there are too many slices to be used in a single day or by allowing better "organ sharing", i.e. allowing the entire brain to be used by several researchers for different experiments, using different areas in separate slices. Moreover, it could reduce preparation efforts for electrophysiological studies and reduce material cost.

This procedure is of very limited use, when the amount of available brain tissue is the limiting factor: when investigating a small brain area or even a subset of neurons that renders merely one or two slices, then these are easily used up in a single day and there is no need for extending viability. Moreover, it is obviously not as successful in areas that are subject to mechanical trauma during preparation, such as L1 neurons of M1/M2 [124], or areas that have dense fibre projections passing through them. However, for experimental procedures that require more time than is typically available from conventional brain slice preparations, such as staining with choleratoxin-B conjugated dyes [131,132], this procedure may make the difference between an *in vivo* experiment, requiring intracranial injections into anaesthetized animals, and an *in vitro/ex vivo* experiment requiring incubated brain slices. Whether, hypothermia of brain slices hampers such tracing experiments remains to be tested.

In conclusion, while this approach may be desirable for reasons of simplifying experimental procedures, reducing cost, and for reasons of animal welfare, it is all the more important that it undergoes independent critical assessment from other scientists of the field in order to become an established procedure. We therefore wish to encourage readers to further test this procedure for flaws, weaknesses, or objectionable alterations to neuronal properties.

## Acknowledgements

The work reported of herein was supported by a grant from the German Research Foundation (DFG) to PC (CO 1334, project number 523878561) and by the Center of Excellence BrainLinks-BrainTools (EXC 1086). The authors wish to thank Christine Zeschnigk, Concepcion Sainz-Rueda, Claudia Roderigo, and Simone Durry for excellent technical assistance. The "UKM" chamber is a joint development of Thomas Budde, Philippe Coulon, Tilman Broicher, Hans-Christian Pape, Manfred Daweke, Ludger Sasse, and Thomas Westhoff and was manufactured by "Fein- und Elektromechanische Forschungswerkstätten, Universitätsklinikum Münster (UKM), Robert-Koch-Str. 27a, 48149 Münster, Germany".

